# Unraveling protein conformational plasticity with PROTEUS

**DOI:** 10.64898/2026.04.27.721098

**Authors:** Luiz Felipe Piochi, Yasaman Karami, Hamed Khakzad

## Abstract

Protein conformational plasticity underpins allosteric regulation, fold switching, and post-translational modification accessibility, yet no existing method can probe this property at the proteome scale without simulation. Here we show that SimpleFold, a flow-matching protein structure predictor, implicitly encodes conformational plasticity in its internal representations. By comparing per-residue embeddings between the sequence-only regime and the structure-converged regime of the denoising trajectory, we define a zero-shot conformational plasticity score, PROTEUS (PROtein TrajEctory Uncertainty Score), that requires no experimental dynamics data. PROTEUS correctly orders five independent protein classes spanning the full flexibility spectrum, from rigid *de novo* designed scaffolds to intrinsically disordered proteins that fold upon binding. Per-residue PROTEUS profiles correlate with atomic fluctuations from 1,290 independent molecular dynamics trajectories, and this signal persists after controlling for structure prediction confidence (pLDDT) and sequence-based disorder predictions. At the protein level, PROTEUS achieves AUROC = 0.77 for fold-switch detection, 0.81 for open/closed state discrimination, and 0.93 for identifying proteins with buried phosphorylation sites. Proteome-wide analysis of 4,188 *Escherichia coli* K-12 proteins reveals that fimbrial adhesins and the type II secretion machinery rank among the most conformationally plastic functional classes, consistent with the structural demands of chaperone-mediated secretion and receptor engagement, while ribosomal proteins score systematically lower. These results establish that PROTEUS provides unsupervised, proteome-scale probing of structural dynamics directly from a generative model.

## 1 Introduction

Proteins are not static objects but dynamic ensembles that sample alternative conformations across timescales ranging from picoseconds to seconds. This conformational plasticity underlies diverse biological phenomena, including allosteric signal transduction, fold switching between structurally distinct states, intrinsic disorder, pathogenicity, and the regulation of enzyme activity and post-translational modification accessibility [1, 2]. Understanding which proteins are conformationally flexible, and to what extent, is therefore a central problem in structural biology and drug discovery. However, direct experimental characterization of conformational dynamics remains challenging. Approaches such as molecular dynamics (MD) simulations, determination of multiple structural states by X-ray crystallography or cryo-EM, or NMR measurements are resource-intensive and difficult to scale to entire proteomes.

Recent advances in deep learning have transformed protein structure prediction. AlphaFold2 [3] demonstrated that neural networks can infer accurate three-dimensional structures directly from sequence. In parallel, large protein language models (pLMs), such as ESM-2 [4] and structure-aware variants like SaProt [5], have shown that latent representations learned from sequence and structure encode rich biological information predictive of protein-protein interactions [6–8], function [9], stability, and mutational effects [10]. Yet these models are fundamentally trained on static representations of proteins and primarily capture evolutionary constraints rather than a protein’s propensity to explore multiple structural states. In reality, proteins occupy a continuum of conformational behaviours. At one extreme are *de novo* designed protein scaffolds engineered to adopt a single rigid fold. Globular proteins typically display modest breathing motions around a dominant structure. Some proteins switch between two or more well-defined folds, while intrinsically disordered proteins (IDPs) populate highly heterogeneous ensembles without a unique native state [11] (**Figure 1A**). Capturing this spectrum of behaviour requires information about the degeneracy of the underlying conformational landscape, which is not directly accessible from evolutionary sequence statistics alone.

**Figure 1:**
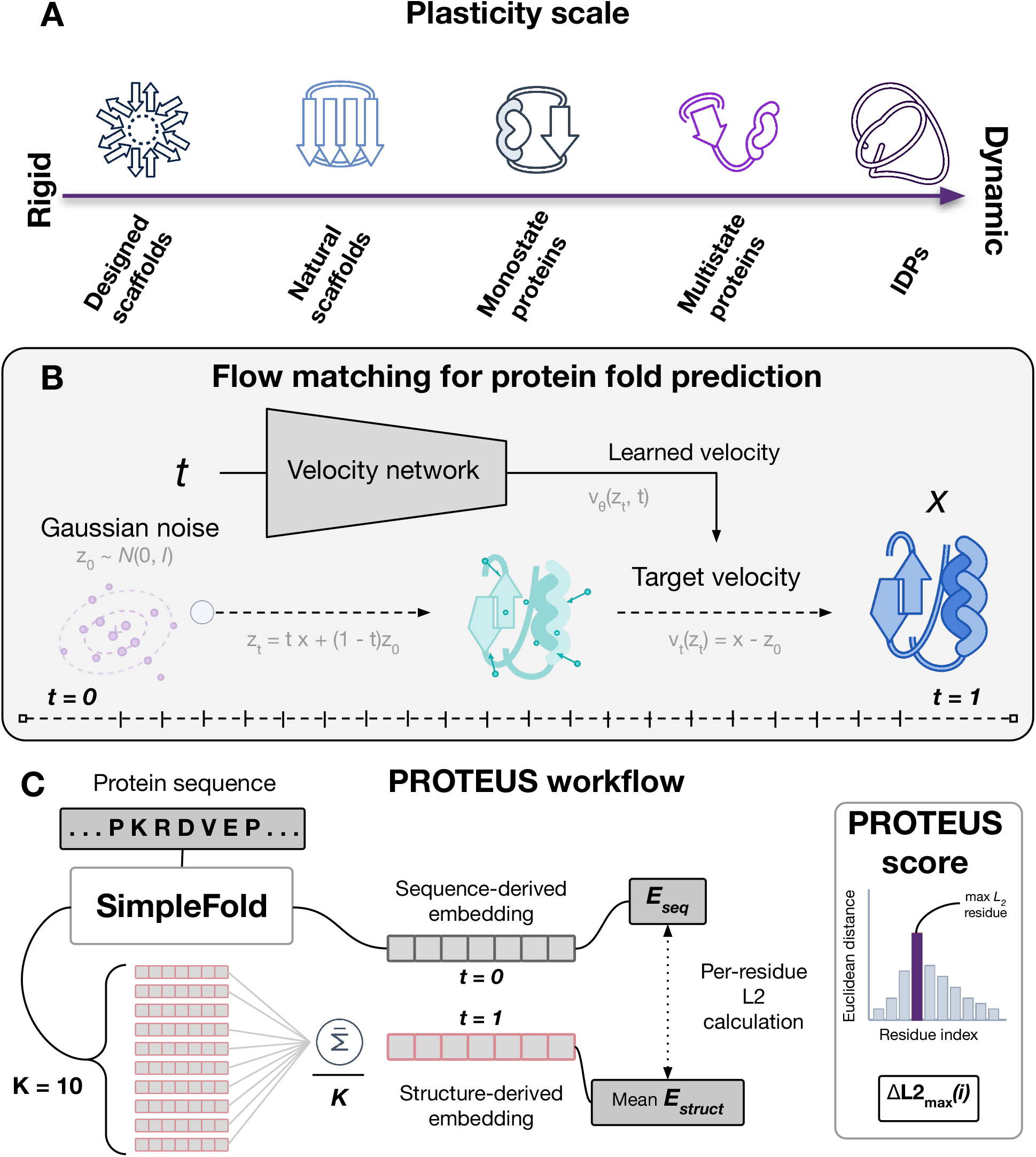
PROTEUS conceptual overview. **A**, Protein conformational plasticity scale illustrating the spectrum from rigid *de novo* designed scaffolds (left) to folding-upon-binding IDPs (right), with schematic icons representing the five protein classes used throughout this study. **B**, Schematic of protein structure prediction using flow-matching strategies. A velocity network is trained to approximate the target velocity field *v*_*t*_(*z*_*t*_) = *x* − *z*_0_ that transports Gaussian noise (*z*_0_ ∼ 𝒩 (0, **I**)) to the distribution of native protein structures. During inference, the Euler-Maruyama sampler follows the learned velocity *v*_*θ*_(*z*_*t*_, *t*) over a discrete trajectory from *t* = 0 (noised) to *t* = 1 (converged folded structure). **C**, PROTEUS scoring workflow. A protein sequence is passed to SimpleFold twice: once with zero coordinates to yield the sequence-only embedding *E*_seq_ at *t* = 0, and *K* = 10 times through the full denoising trajectory to yield structural embeddings averaged into the mean structural embedding *Ē*_struct_ at *t* = 1. The PROTEUS score, Δ*L*2_max_(*i*), is the maximum per-residue Euclidean distance between *E*_seq_ and *Ē*_struct_, quantifying the largest embedding displacement induced by introducing structural information.

Flow-matching generative models offer an alternative perspective on protein structure prediction [12, 13]. Rather than producing a single deterministic structure, these models learn a continuous velocity field that transports samples from a noise distribution toward the distribution of native protein structures conditioned on sequence [14]. During inference, coordinates are iteratively refined through a denoising trajectory, typically using an Euler-Maruyama sampler. Crucially, the information available to the model changes along this trajectory. At early timesteps, coordinates are close to pure noise and the model’s internal representation is dominated by sequence information, effectively reflecting the ensemble of structures compatible with the sequence. At late timesteps, when the sampler has converged, the representation corresponds to a specific three-dimensional conformation. These two regimes therefore capture distinct aspects of the sequence-structure relationship: the sequence-constrained structural prior and the final realized fold (**Figure 1B**).

We leverage this framework using *SimpleFold*, a recently introduced flow-matching structure predictor, which allows access to internal representations throughout the denoising trajectory [15]. We hypothesize that the displacement between these two regimes in representation space provides a proxy for conformational flexibility (**Figure 1C**). For proteins with a single well-defined fold, the sequence strongly constrains the final structure, and the model representation changes only modestly as structural information is introduced. In contrast, sequences compatible with multiple structural states, such as fold-switching proteins or IDPs, should exhibit larger shifts in representation space as the denoising trajectory commits to one particular conformation. Conceptually, this interpretation aligns with recent arguments that diffusion and flow-matching models implicitly encode aspects of the conformational landscape through their learned transport dynamics [16].

We operationalize this idea with PROTEUS (**PRO**tein **T**raj**E**ctory **U**ncertainty **S**core). Given a protein sequence, PROTEUS extracts residue-level embeddings from SimpleFold at two points along the denoising trajectory: a sequence-only regime, obtained by removing coordinate information, and a structure-converged regime, obtained after running the full Euler-Maruyama sampling trajectory. The maximum residue-level Euclidean (L2) distance between these embeddings defines the PROTEUS score. Importantly, PROTEUS is a zero-shot approach that bypasses a major limitation of supervised flexibility predictors, namely the scarcity of large experimental datasets describing protein dynamics. We first demonstrate that PROTEUS correctly orders five independent protein classes spanning the conformational flexibility spectrum, from *de novo* designed rigid scaffolds to intrinsically disordered proteins that fold upon binding. We then establish the mechanistic basis of the signal by validating against per-residue root-mean-square fluctuations derived from 1,290 molecular dynamics trajectories, showing that the correlation persists after controlling for pLDDT and disorder prediction. Furthermore, we show that the same score detects open-closed conformational transitions and identifies proteins containing buried phosphorylation sites. Finally, we apply PROTEUS at proteome scale to *E. coli*, revealing a systematic association between predicted conformational plasticity and functional categories involved in host interaction and protein secretion.

## 2 Results

### 2.1 PROTEUS quantifies conformational plasticity from flow-matching trajectories

We built PROTEUS on top of SimpleFold [15], a protein structure predictor trained on the protein data bank (PDB) via a generative flow-matching objective. SimpleFold uses an Euler-Maruyama sampler running for *n* steps over a uniform flow schedule from pure noise towards a converged structure. At each step, the model receives noised coordinates together with sequence features and outputs a predicted velocity vector. The internal trunk representation (which we refer to as the latent embedding) has shape *L* × *D*, where *L* is protein sequence length and *D* = 1024.

We extract embeddings at two regimes (**Figure 1C**). The sequence-only embedding is obtained by a single forward pass with coordinates set to **0**, corresponding to flow timestep *t* = 0, so that the model receives no structural information. The structural ensemble is obtained by running the sampler *K* = 10 times from independently sampled noise vectors, with *K* chosen by ablation experiments (Supplementary Figure S1), and the structural embedding is the mean of the *K* per-residue latent representations at the end of the trajectory (*t* = 1).

The primary conformational plasticity score is defined as

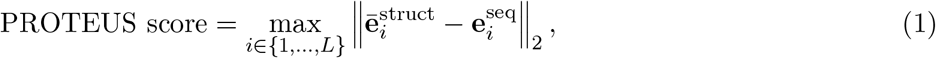

where 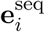 is the sequence-only embedding at residue *i* and 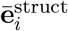 is the mean structural embedding at residue *i* over the *K* conformational samples. Taking the maximum rather than the mean captures the most ambiguous residue, which we find empirically to be more discriminative than whole-protein averages (see Supplementary Figure S2). In other words, the PROTEUS score is the maximum per-residue Euclidean displacement between the sequence-only embedding and the mean structure-conditioned embedding, quantifying how much the model’s representation of the most flexible residue shifts as structural information is added.

To select the best formulation, we screened approximately 70 candidate scoring functions derived from the same trajectory embeddings, including per-residue L2 norms, cosine distances, ensemble spread (variance across conformational samples), and trajectory nonlinearity metrics, evaluated on the fold-switch benchmark. The final PROTEUS score (l2 delta max) consistently ranked first or tied for first across folds (**Supplementary Figure S3**). Trajectory nonlinearity (the deviation of latent embeddings at *t* = 0.5 from the linear interpolation between endpoints) achieved AUROC values in the range 0.36–0.40, confirming that the signal is concentrated at the endpoints of the trajectory rather than in intermediate curvature.

### 2.2 PROTEUS scores capture the continuum of conformational plasticity

We further asked if PROTEUS captures a genuine property of protein conformational plasticity. If so, its scores should correctly order protein classes whose flexibility is independently established. We assembled five such classes and scored each with the PROTEUS score, residualising scores on sequence length to remove the trivial dependence of the maximum on protein size (**Figure 2A**).

**Figure 2:**
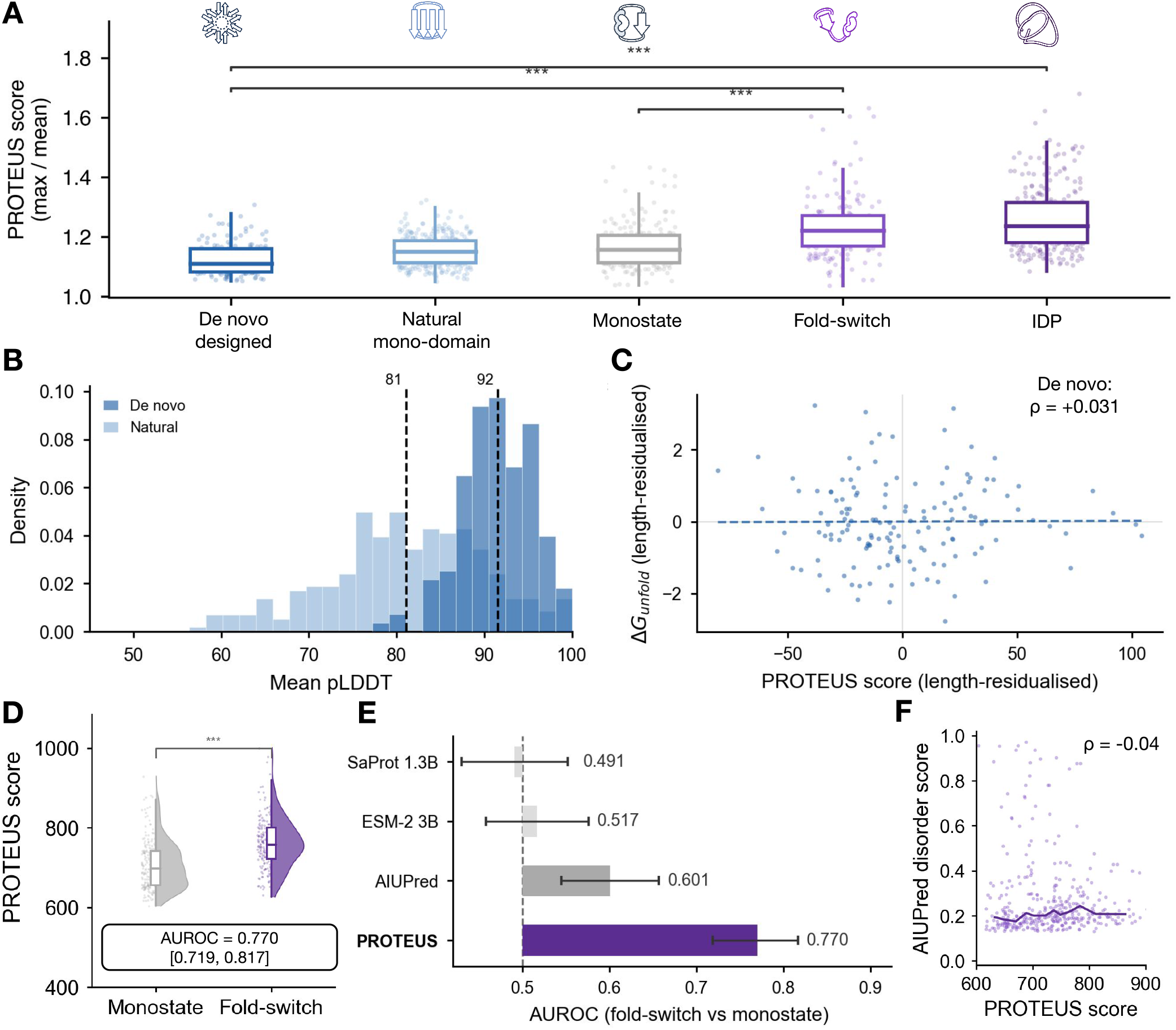
PROTEUS scores capture the continuum of protein conformational plasticity. **A**, Max-mean-normalized PROTEUS scores for five independent protein classes spanning the full conformational flexibility spectrum: *de novo* designed miniproteins and natural single-domain proteins (Tsuboyama 2023 dataset), monostate proteins and fold-switching proteins (fold-switch benchmark), and intrinsically disordered proteins (DIBS database). Significant pairwise differences are annotated (Mann-Whitney U test, *** *p <* 0.001). **B**, Distribution of mean pLDDT for *de novo* designed (dark blue) and natural single-domain (light blue) proteins from the Tsuboyama dataset. Dashed lines indicate median values 81 and 92, respectively, confirming that *de novo* proteins are structurally well-defined. **C**, Scatter plot of length-residualised PROTEUS score versus experimentally measured unfolding free energy (Δ*G*_unfold_) for *de novo* designed proteins. Partial Spearman *ρ* = 0.031 indicates no association between thermodynamic stability and PROTEUS score within this class. **D**, PROTEUS score distributions for monostate (*n* = 197) and fold-switching (*n* = 189) proteins shown as half-violin plots with individual data points. Fold-switching proteins score significantly higher (AUROC = 0.770 [0.719, 0.817], *** *p <* 0.001, Mann-Whitney *U* test). **E**, AUROC for zero-shot fold-switch versus monostate classification for PROTEUS and AIUPred, ESM-2 3B, and SaProt 1.3B. Error bars indicate bootstrap 95% confidence intervals. A dashed line indicates chance level (AUROC = 0.5). **F**, Scatter plot of maximum per-residue AIUPred disorder score versus PROTEUS score for fold-switching proteins (*ρ* = − 0.04), demonstrating that PROTEUS scores are orthogonal to sequence-based disorder tendency within the fold-switch class.

*De novo* designed proteins are explicitly engineered to adopt a unique, highly stable fold with minimal conformational degeneracy. We extracted wild-type sequences from the Tsuboyama 2023 mega-scale stability dataset [17], comprising 146 de novo designed miniproteins and 308 natural proteins (Dataset 2, lengths 44 or 72 amino acids after library padding), and scored them with PROTEUS. *De novo* designed proteins obtained the lowest mean length-residualised PROTEUS score of all five classes, below natural single-domain proteins from the same dataset. Because these proteins are well-folded (high pLDDT by construction), their low PROTEUS score cannot be attributed to disorder; rather, it reflects the absence of alternative structural states in their conformational landscape. Within the Tsuboyama set, length-controlled partial Spearman correlation between PROTEUS and experimentally measured folding free energy Δ*G* was negligible for *de novo* proteins (*ρ* = 0.031, *p* = 0.71), indicating that variation in stability within the *de novo* class does not further stratify the PROTEUS score once sequence length is accounted for.

Monostate proteins from the fold-switch benchmark [18] (*n* = 197) scored intermediate between natural single-domain proteins and fold-switching proteins (*n* = 189), with the fold-switch group scoring significantly higher (Mann-Whitney *p* = 1.6×10^−15^). Quantitatively, PROTEUS achieved AUROC = 0.770 [0.719–0.817] (bootstrap 95% confidence interval) for zero-shot fold-switch detection (**Figures 2D,E**), with a length-matched Wilcoxon signed-rank test confirming the separation is not driven by the length difference between groups (*p* = 1.3 × 10^−13^). A permutation test (5,000 label shuffles) yielded *p* = 0. Partial Spearman *ρ* after jointly controlling for sequence length and crystallographic resolution was 0.161 (*p* = 4.4 × 10^−3^).

By contrast, ESM-2 3B [4] and SaProt 1.3B [5] population-deviation scores both achieved AUROC values indistinguishable from chance (ESM-2 AUROC = 0.50, SaProt AUROC = 0.50, **Figure 2E**). AIUPred [19], an AI-based intrinsic disorder predictor, achieved AUROC = 0.601 for its maximum per-residue disorder score after length control (partial *ρ* = 0.179), substantially below PROTEUS. These results confirm that trajectory displacement encodes qualitatively different information from evolutionary statistics or disorder tendency.

Proteins from the DIBS database [20], which catalogues intrinsically disordered regions that undergo disorder-to-order transitions upon binding to a structured partner (folding-upon-binding), obtained the highest mean length-residualised PROTEUS score for all five classes. The separation from fold-switching proteins was highly significant (Mann-Whitney *p* = 2.5 × 10^−24^), and from monostate proteins even more so (*p* = 3.6 × 10^−53^).

Taken together, the five classes establish a monotone ordering of the PROTEUS score: *de novo* designed *<* natural single-domain *<* monostate *<* fold-switching *<* DIBS folding-upon-binding (**Figure 2A**). Non-adjacent and boundary comparisons involving qualitatively different flexibility classes are highly significant (*de novo* vs fold-switch: *p* = 2.2 × 10^−25^ and *de novo* vs DIBS: *p* = 7.5 × 10^−56^), while the natural/monostate adjacent comparison is not significant in isolation because the two classes differ substantially in length (Tsuboyama proteins: 44–72 aa; monostate benchmark: 17–510 aa) and span overlapping conformational regimes. This ordering is consistent with PROTEUS capturing a genuine gradient of conformational plasticity rather than a binary switch.

### 2.3 Per-residue trajectory displacement correlates with MD fluctuations

The continuum results demonstrate that PROTEUS correctly orders protein classes at the protein level. We next asked whether the per-residue PROTEUS profile quantitatively reflects the actual atomic fluctuations of individual residues as measured by molecular dynamics.

For each of 1,290 proteins from the ATLAS database [21], we computed the within-protein Spearman correlation between the per-residue PROTEUS profile and the mean per-residue RMSF from three independent 100 ns molecular dynamics trajectories. Representative proteins coloured by RMSF and PROTEUS score show good spatial concordance (**Figures 3A,B**; extended galleries in **Supplementary Figures S5** and **S6**). The median within-protein correlation was 0.396 (mean 0.382; **Figure 3C**). This result holds across all secondary-structure classes: all-alpha proteins (median *ρ* = 0.420), all-beta proteins (median *ρ* = 0.395), and mixed proteins (median *ρ* = 0.378), ruling out secondary-structure composition as a confound. At the protein level, PROTEUS correlated with mean RMSF across all 1,290 proteins at partial Spearman *ρ* = 0.403 (*p <* 10^−30^) after controlling for sequence length.

**Figure 3:**
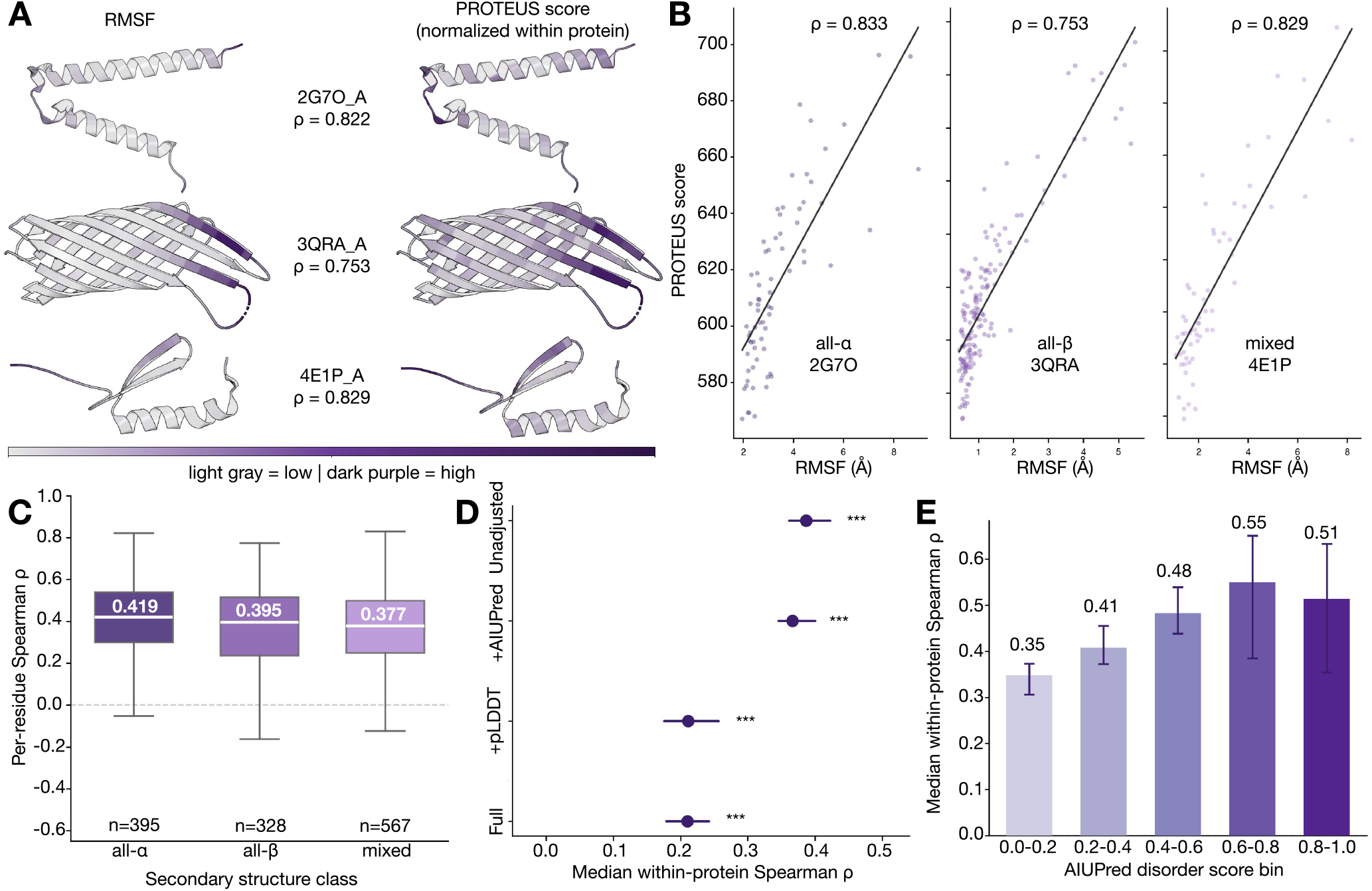
Per-residue PROTEUS profiles correlate with molecular dynamics fluctuations. **A**, Three representative proteins from the ATLAS database coloured by mean MD RMSF (left) and min-max normalised PROTEUS score (right): an all-*α* protein (2G7O_A, *ρ* = 0.822), an all-*β* protein (3QRA_A, *ρ* = 0.753), and a mixed secondary-structure protein (4E1P_A, *ρ* = 0.829). Colour scale from light grey (low) to dark purple (high). **B**, Scatter plots of per-residue PROTEUS score versus mean MD RMSF for the same three proteins. Each point represents one residue; Spearman *ρ* values are shown. **C**, Distribution of within-protein Spearman *ρ* between per-residue PROTEUS score and mean MD RMSF, stratified by SCOP secondary structure class (all-*α*: *n* = 395, median *ρ* = 0.419; all-*β*: *n* = 328, median *ρ* = 0.395; mixed: *n* = 567, median *ρ* = 0.377). Box plots show median, interquartile range, and 1.5 × IQR whiskers. A dashed line is shown at *ρ* = 0. **D**, Median within-protein Spearman *ρ* between PROTEUS and MD RMSF under successive confound control, evaluated on a stratified subset of 300 ATLAS proteins: unadjusted, controlling for AIUPred disorder score alone (+AIUPred), controlling for per-residue pLDDT alone (+pLDDT), and jointly controlling for both (Full). Error bars indicate bootstrap 95% confidence intervals, *** *p <* 0.001 against the null hypothesis of zero correlation. **E**, Median within-protein Spearman *ρ* between PROTEUS and MD RMSF stratified by AIUPred disorder score bin, from fully ordered residues (0.0–0.2, *ρ* = 0.35) to highly disordered residues (0.8–1.0, *ρ* = 0.51). Error bars indicate bootstrap 95% confidence intervals. The PROTEUS–RMSF correlation is positive across all disorder bins, demonstrating that the signal is not restricted to disordered residues.

Both pLDDT and sequence-based disorder predictions are known correlates of RMSF. To determine how much of the PROTEUS signal is independent of these confounds, we ran AIUPred [19] on a stratified subset of 300 ATLAS proteins, selected to span the full range of sequence length, mean RMSF, and structural class represented in the database, and computed partial Spearman correlations between the PROTEUS score and RMSF after successive confound controls (**Figure 3D**).

Controlling for AIUPred disorder alone reduced the median within-protein correlation only marginally (from 0.387 to 0.366), demonstrating that sequence-based disorder prediction and PROTEUS carry nearly non-overlapping per-residue information. Controlling for per-residue pLDDT reduced the correlation more substantially (from 0.387 to 0.211), consistent with pLDDT and PROTEUS both reflecting the degree to which a residue is structurally committed (**Supplementary Figure S4**). Critically, jointly controlling for both pLDDT and AIUPred yielded an identical residual correlation (0.210), confirming that the AIUPred confound is fully subsumed by pLDDT and that PROTEUS retains an independent signal of ≈ 0.21 after all available confound control. Notably, pLDDT and AIUPred themselves showed only modest per-residue correlation (median within-protein *ρ* = 0.257), confirming they are distinct confounds and that controlling for one does not substitute for the other.

If PROTEUS merely detected disordered residues, its correlation with RMSF should vanish when restricted to ordered residues. Restricting to residues with AIUPred score *<* 0.2 to retain the most ordered residues by sequence-based prediction still yielded a positive median within-protein correlation with RMSF (*ρ* ≈ 0.35 for *n* = 279 proteins with sufficient ordered residues) (**Figure 3E**). Across AIUPred bins from 0.0–0.2 (fully ordered) to 0.8–1.0 (highly disordered), the PROTEUS-RMSF correlation increased from 0.35 through 0.41, 0.53, and 0.59 for successively more disordered bins, remaining robustly positive throughout (**Figure 3E**). This establishes that PROTEUS captures structured conformational plasticity and is not acting merely as a proxy for disorder.

### 2.4 PROTEUS detects biologically relevant conformational plasticity

We next evaluated the utility of PROTEUS score for detecting specific biological manifestations of conformational plasticity. To test whether PROTEUS detects open/closed conformational switching, we used two complementary benchmarks that differ in the magnitude of the structural change. The OC23 dataset [22] comprises 23 proteins each crystallised in a functionally open (apo or substrate-accessible) and a functionally closed (holo or occluded) conformation, with TM-score *<* 0.85 between the two states. The OC85 dataset [22] comprises 21 proteins from the same experimental paradigm but with TM-score *>* 0.85 between open and closed states. For both benchmarks, we compared PROTEUS scores against two negative controls: the 197 monostate proteins from the fold-switch benchmark and 380 MD-validated rigid proteins from ATLAS (mean RMSF ≤ 1.0 Å, a positive experimental measure of rigidity).

OC23 proteins scored significantly higher than both controls (**Figure 4A**). Against monostate proteins PROTEUS achieves an AUROC = 0.808 (partial *ρ* = 0.209; **Figure 4B**). Against ATLAS rigid proteins it achieves AUROC = 0.776 (partial *ρ* = 0.071, *p* = 0.16). The partial *ρ* becomes non-significant against this control, reflecting the limited statistical power with *n* = 23.

**Figure 4:**
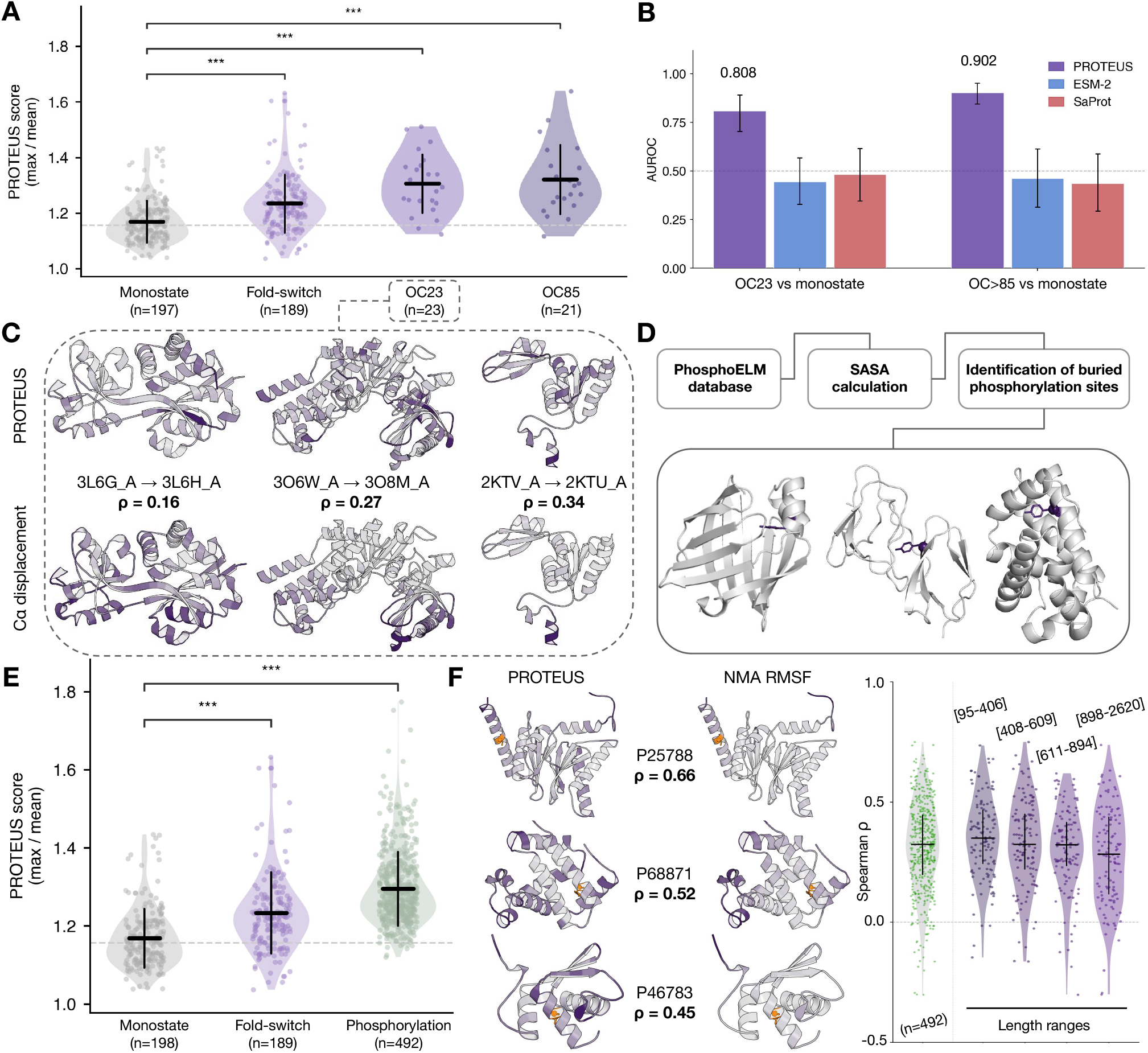
PROTEUS detects biologically relevant conformational plasticity. **A**, PROTEUS score distributions for monostate proteins (*n* = 197), fold-switching proteins (*n* = 189), and proteins with crystal-lographically observed open/closed conformational states with TM-score *<* 0.85 (OC23, *n* = 23) or TM-score *>* 0.85 (OC85, *n* = 21) between states. Violin plots show the full distribution; + marks indicate the median. Significance brackets indicate Mann-Whitney *U* test results (*** *p <* 0.001). A dashed line indicates the monostate median. **B**, AUROC for zero-shot discrimination of open/closed proteins against monostate controls for PROTEUS (purple), ESM-2 3B (blue), and SaProt 1.3B (red), evaluated on the OC23 and OC85 benchmarks. Error bars indicate bootstrap 95% confidence intervals. A dashed line indicates chance level (AUROC = 0.5). **C**, Three representative open/closed protein pairs from the OC23 dataset coloured by per-residue PROTEUS score (top row) and C*α* displacement between open and closed states (bottom row). Spearman *ρ* values indicate within-protein correlation between the PROTEUS residue profile and the observed C*α* displacement. **D**, The pipeline of buried phosphorylation site dataset from PhosphoELM, with three examples shown below. **E**, PROTEUS score distributions for monostate, fold-switch, and phosphorylation-site proteins (*n* = 492). Proteins with buried phosphorylation sites score significantly higher than monostate controls (*** *p <* 0.001, Mann-Whitney *U* test). **F**, Left: three representative phosphorylation-site proteins (P25788, P68871, P46783) coloured by per-residue PROTEUS score (left) and NMA RMSF (right); buried phosphorylation sites highlighted in orange. Right: distribution of within-protein Spearman *ρ* between PROTEUS and NMA RMSF across four protein length ranges within the full phosphorylation dataset (*n* = 492).

OC85 proteins, despite representing only subtle structural transitions, scored higher than OC23 proteins on PROTEUS and achieved stronger discrimination against both controls (**Figure 4A**). Against monostate: AUROC = 0.902 (partial *ρ* = 0.273; **Figure 4B**). Against ATLAS rigid proteins: AUROC = 0.872 (partial *ρ* = 0.189). The partial *ρ* for OC85 survives length control against the more stringent ATLAS negative set, whereas OC23 does not. This counter-intuitive result suggests that PROTEUS detects the conformational plasticity of a sequence rather than the magnitude of its largest observed structural excursion. Given the small sample sizes, larger open/closed benchmarks would be needed to confirm these trends.

Importantly, ESM-2 3B population-deviation scores on OC23 yielded AUROC = 0.321 (**Figure 4B**), meaning ESM-2 actively assigns lower scores to open/closed proteins than to monostate controls, suggesting that the signal captured by PROTEUS is not accessible to static sequence representations.

We next assembled a dataset of 492 human proteins harbouring at least one experimentally validated buried phosphorylation site from PhosphoELM [23] (see Methods section for more details). Phosphorylation is a ubiquitous regulatory mechanism that modifies serine, threonine, and tyrosine residues to alter protein activity, localization, and interaction networks. In some cases, the regulatory phosphorylation events target residues buried in the protein core, which can only be phosphorylated if the protein transiently samples conformations that expose those residues to a kinase active site. We therefore, hypothesized that the existence of a buried phosphorylation site implies that the host protein must be conformationally flexible enough to expose it.

We compared the proteins with buried phosphorylation sites against both the 197 monostate proteins from the fold-switch benchmark and 380 MD-validated rigid ATLAS proteins (mean RMSF ≤ 1.0 Å) (**Figure 4E**). Proteins harbouring buried phosphorylation sites scored substantially higher on all PROTEUS features. Against monostate proteins PROTEUS scores an AUROC = 0.930 (partial *ρ* = 0.188), whereas against ATLAS rigid proteins it achieves AUROC = 0.914 (partial *ρ* = 0.285).

In contrast, site-level PROTEUS scores did not achieve significant discrimination compared with unphosphorylated buried residues of the same protein, consistent with the signal residing in whole-protein conformational flexibility rather than local residue-level variability. The protein-level result alone, however, provides a biologically interpretable prediction that proteins harbouring buried phosphorylation sites are globally more flexible, and PROTEUS detects this without any training on modification data. Given that the PROTEUS score correlates with per-residue RMSF from ATLAS but cannot discriminate phosphorylated from unphosphorylated buried residues within the same protein, we infer that the signal is driven by global conformational plasticity rather than local flexibility of the phosphorylation residue/site itself. An extended gallery of buried-phosphorylation-site proteins coloured by per-residue PROTEUS score is provided in **Supplementary Figure S7**.

### 2.5 Proteome-wide mapping of conformational plasticity in *Escherichia coli* K-12

As a proteome-scale application, we applied PROTEUS to the complete reference proteome of *Escherichia coli* K-12 (UP000000625), comprising 4,188 proteins in the length range 50–2000 amino acids. We filtered out disordered proteins using AIUPred (mean AIUPred ≤ 0.3, fraction of disordered residues ≤ 0.3) to focus on structured proteins for which PROTEUS is most interpretable, retaining 3,549 proteins (84.7%) as ordered candidates for analysis. Functional annotation of the 3,549 scored proteins reveals that the highest PROTEUS scores are concentrated in specific functional niches rather than distributed uniformly across the proteome (**Figure 5A**). Fimbrial adhesins and secretion system components carry the highest median scores, followed by ABC transporters and chaperones, while ribosomal proteins are the most depleted category among high scorers (**Figure 5B**). This ordering is biologically coherent. Fimbrial tip adhesins and type II/III secretion needle components must cycle between retracted and extended states during host contact or substrate translocation, whereas the ribosome operates as a conformationally constrained molecular machine whose core rRNA-protein contacts are under strong evolutionary pressure for rigidity.

**Figure 5:**
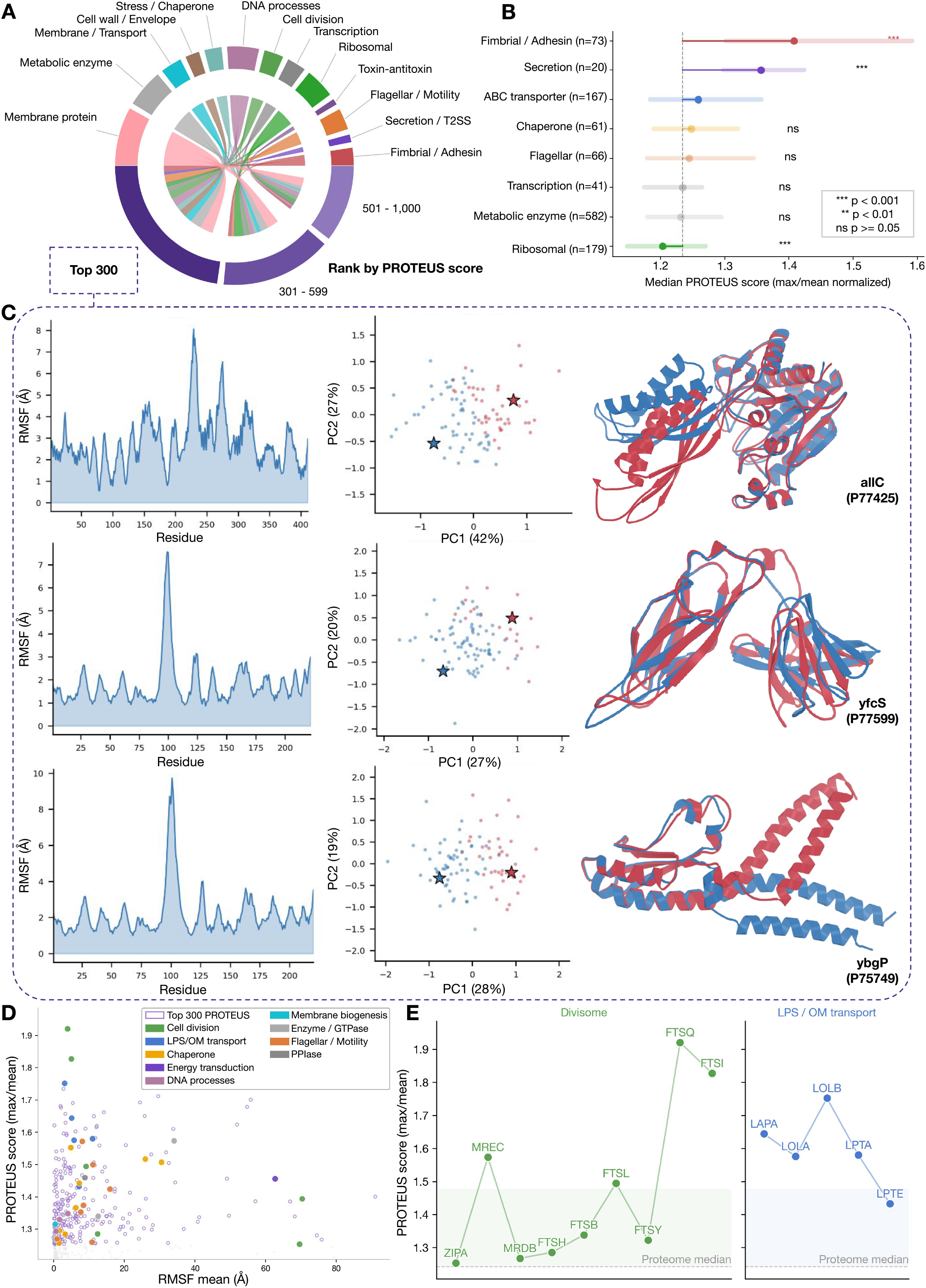
Proteome-wide conformational plasticity landscape of *Escherichia coli* K-12. **A**, Circular sunburst chart of the 4,188 scored *E. coli* K-12 proteins distributed by broad functional category (inner ring) and specific subcategory (outer ring), ranked by PROTEUS score. The top 300 highest-scoring proteins are highlighted. **B**, Median PROTEUS score (max/mean normalised) for selected functional categories relative to the proteome-wide median (dashed vertical line). Horizontal error bars indicate bootstrap 95% confidence intervals. Statistical significance against the proteome background: *** *p <* 0.001, ** *p <* 0.01, ns *p* ≥ 0.05 (Mann-Whitney *U* test). **C**, Detailed characterisation of three high-ranking uncharacterised proteins from the top 300: allC (P77425), yfcS (P77599), and ybgP (P75749). For each protein: right, two states (one in blue, and one in red) from different conformations sampled using BBFlow; centre, PCA of all sampled conformations by pairwise RMSD comparisons; left, mean RMSF per residue from ANM analysis. **D**, Scatter plot of PROTEUS score versus mean ANM RMSF for all proteome, with top-300 proteins shown as filled circles coloured by functional annotation and remaining proteins shown as open circles. **E**, PROTEUS scores for individual members of the divisome pathway (left) and the LPS/outer-membrane (LPS/OM) transport pathway (right), with the proteome-wide median indicated by a dashed horizontal line. The colored background area represents the proteome 90th percentile.

To characterise the conformational diversity of the top 300 highest-scoring proteins, we generated structural ensembles using BBFlow [24]. Three functionally uncharacterised proteins, allC (P77425), yfcS (P77599), and ybgP (P75749), serve as illustrative examples (**Figure 5C**). For allC and yfcS, principal component analysis of pairwise RMSD between sampled conformations reveals two clearly separated clusters in PC space, with the query structure (dark star) offset from the ensemble 10th/90th quantiles, indicative of two distinct conformational states rather than diffuse local fluctuations. The corresponding anisotropic network model (ANM) RMSF profiles show broad, multi-residue peaks consistent with domain-level rearrangements. By contrast, ybgP displays a single sharp RMSF peak localised around residue ∼100, suggesting a discrete hinge that connects two otherwise rigid subdomains. In all three cases, the PROTEUS score correctly anticipates the conformational heterogeneity revealed by explicit ensemble sampling, providing functional hypotheses for proteins that lack experimental characterisation. ANM RMSF, BBFlow PCA, and ensemble-overlay triplets for the top 50 *E. coli* hits are shown in **Supplementary Figure S8**.

Plotting PROTEUS score against mean ANM RMSF for all 3,549 proteins exposes a positive but non-linear relationship (**Figure 5D**). Multiple functionally coherent clusters emerge in the top-300 subset, with membrane biogenesis, cell division, and LPS/OM transport proteins consistently occupying the high-score, high-RMSF quadrant. Among the most striking results is the systematic elevation of PROTEUS scores across entire functional pathways (**Figure 5E**). All scored members of the divisome, including FtsQ, FtsL, FtsB, FtsI, FtsH, MreC, and ZipA, score above the proteome median, and several exceed the 90th percentile. The FtsQ/FtsL/FtsB sub-complex is known to act as a conformational switch that couples membrane tension sensing to activation of the cell-wall synthesis machinery [25, 26]. A similar picture holds for the LPS/outer-membrane transport system, where LptA, LolA, LolB, LptE, and LptD all score above the proteome median [27, 28]. LptA forms a periplasmic bridge by adopting extended oligomeric conformations that must open to receive LPS from the inner membrane and close to pass it along the chain, while LolA and LolB require a hydrophobic cavity opening to bind and release lipoprotein cargo. Taken together, these results suggest that the PROTEUS score reflects function-related conformational plasticity in proteome-level calculations.

## 3 Discussion

In this study, we have shown that the displacement between sequence-only and structure-conditioned embeddings along the denoising trajectory of SimpleFold serves as a robust zero-shot proxy for protein conformational plasticity across the full flexibility spectrum. The primary score, l2 delta max, correctly orders five independent protein classes from rigid *de novo* designed scaffolds to folding-upon-binding IDPs, correlates with per-residue atomic fluctuations in molecular dynamics at residue resolution, and detects biologically relevant conformational events including domain opening, fold switching, and buried post-translational modification site accessibility. Static protein language models and disorder predictors fail on these same tasks, confirming that trajectory-based representations encode a qualitatively different form of structural information.

The PROTEUS score also provides a principled, zero-shot measure on conformational flexibility that spans the full spectrum of protein design objectives, from the engineering of maximally rigid scaffolds to the ambitious goal of designing multi-state proteins that switch folds on demand. In de novo design, where methods such as RFdiffusion [29] and ProteinMPNN [30] optimise for a single target conformation, the central quality criterion is conformational singularity, meaning that the designed sequence commits to a unique structural state. This is precisely the condition reflected by a low PROTEUS score. Critically, the absence of any correlation between PROTEUS and thermodynamic stability (Δ*G*) within the de novo class (*ρ* = 0.031, *p* = 0.71) demonstrates that PROTEUS captures a dimension of design quality that energy-based metrics cannot. A designed sequence may be thermodynamically stable yet carry an elevated PROTEUS score, signalling a propensity for alternative conformations that would be invisible to pLDDT or folding-free-energy filters. Incorporating PROTEUS as a post-design screening step thus provides orthogonal quality control for rigidity-critical applications such as biosensors, nanocages, and therapeutic scaffolds. At the opposite end of the design spectrum, where the goal is to engineer functional flexibility, such as flexible active-site loops, hinge motions, or binding-induced conformational changes, the per-residue PROTEUS profile offers residue-level guidance without the cost of molecular dynamics. The median within-protein correlation between the per-residue profile and MD-derived RMSF of 0.396 means that PROTEUS provides a rapid, simulation-free prediction of where a designed sequence will be mobile in solution. This creates a practical route to prescribing flexibility patterns by enforcing low per-residue PROTEUS scores at a structural core while permitting elevated scores at a designed interface or catalytic loop. The most ambitious design challenge, programming fold-switching or multi-state proteins, remains largely unsolved by current methods [31] owing to the difficulty of simultaneously satisfying two structurally dissimilar backbones at the sequence level. PROTEUS is uniquely positioned to assist here, as it detects natural fold-switching proteins with AUROC = 0.770 in a zero-shot setting where static protein language models perform at chance, indicating that the flow-matching trajectory encodes the conformational degeneracy that defines a fold-switcher. Taken together, considering the three design regimes of rigid scaffolds, dynamic proteins, and multi-state design position PROTEUS as a flexibility-aware complement to existing computational design pipelines across the full rigidity to plasticity continuum.

The mechanistic basis of the PROTEUS signal is in fact, consistent with recent theoretical arguments that learned denoising networks implicitly encode aspects of the protein conformational landscape. When a protein sequence is compatible with a single stable fold, the denoising velocity field acts as a contractive mapping that rapidly collapses the coordinate distribution toward a point attractor; the model’s internal representation therefore changes little between the sequence-only and structure-converged regimes. When a sequence is compatible with multiple alternative structures of similar energy, the velocity field must first resolve this degeneracy, and the internal representation undergoes a larger rotation as structural information from the specific sampled conformation overrides the sequence-level prior. This interpretation is supported by the trajectory analysis showing that the signal is front-loaded at the start of the denoising trajectory (the sequence-to-intermediate leg), where the model transitions from the degenerate prior to committing to a specific structural state. The intermediate-to-structure leg contributes negligible additional discrimination, consistent with the coordinate updates at late steps being largely refinements within a single state rather than multi-state events.

A particularly important finding is that the PROTEUS signal persists in fully ordered residues (AIUPred *<* 0.2: *ρ* ≈ 0.35 with RMSF), survives joint control for pLDDT and sequence-based disorder prediction (partial *ρ* = 0.210), and yet is essentially orthogonal to AIUPred disorder prediction alone (controlling for AIUPred changes the correlation by less than 0.02). This dissociation reveals that the signal is not equivalent to disorder detection but captures structured conformational plasticity: the tendency of well-folded residues to sample alternative positions. Importantly, the pLDDT and AIUPred confounds are distinct from each other (median within-protein *ρ* = 0.257), and the PROTEUS residual after joint control (0.210) represents genuine information about conformational plasticity that is not accessible from either existing predictor. The finding that PROTEUS and pLDDT carry partially orthogonal information about MD-derived RMSF has important practical implications. pLDDT is already widely used as a proxy for disorder and flexibility. PROTEUS provides an independent, complementary signal, and the two scores can be combined straight-forwardly. Crucially, the information captured by PROTEUS is not accessible to any existing static pLM, confirming the core claim that SimpleFold encodes a qualitatively new form of structural information by virtue of its trajectory-based inference procedure.

Several limitations deserve acknowledgement. First, SimpleFold is trained on static PDB structures, not on experimental dynamic ensembles. The signal it encodes therefore reflects learned correlations between sequence features and structural diversity in the PDB, rather than a direct simulation of thermodynamic fluctuations. Second, the fold-switch benchmark contains proteins annotated by the binary fold-switch label, which conflates proteins that switch under physiological conditions with those where switching was observed only under non-physiological conditions. Third, the ATLAS RMSF data reflect specific simulation conditions (three 100 ns runs) that may not capture slow conformational dynamics on millisecond timescales. Fourth, PROTEUS was validated exclusively on SimpleFold. The two-regime extraction logic applies in principle to any flow-matching structure predictor that jointly conditions on sequence and coordinates throughout the denoising trajectory, but whether other models encode a comparable conformational signal in their internal representations remains to be established empirically.

Looking forward, several extensions of PROTEUS are straightforward to implement. Per-residue l2 delta profiles could identify specific flexible loops or domains within an otherwise rigid protein. Extension to protein complexes, by conditioning the flow-matching sampler on the full complex sequence while comparing single-chain versus complex-chain embeddings, could identify interface plasticity. Finally, the PROTEUS score could serve as a prior or regulariser in molecular dynamics enhanced-sampling workflows, focusing computational effort on proteins identified as conformationally ambiguous.

Finally, the proteome-scale *E. coli* application illustrates that PROTEUS is practical at scale. Scoring the full 4,188-protein proteome required around 2 GPU hours using the optimised inline-latent-capture pipeline, making proteome-wide screening routine on modern hardware. The enrichment of fimbrial, T2SS, and flagellar proteins among high-scoring candidates, and the depletion of ribosomal proteins, provides an immediately interpretable biological result that the functional categories most associated with conformational plasticity during assembly and secretion score highest on PROTEUS, while the most constrained molecular machines score lowest.

## 4 Author Contributions

LP, YK, and HK conceived the idea. LP developed the method in discussion with YK and HK, and performed the data analysis. HK and YK supervised the project. LP wrote the initial draft. All authors contributed to writing the manuscript.

## 5 Acknowledgements

HK was supported by the French National Research Agency (ANR), under grants ANR-22-CPJ2-0075-01, and ANR-24-CE45-4243-01. YK was supported by the French National Research Agency (ANR) under the France 2030 grant reference number ANR-24-RRII-0002 operated by the Inria Quadrant Program.

## 7 Methods

### 7.1 Datasets

The Tsuboyama 2023 mega-scale stability dataset [17] provided 146 *de novo* designed miniproteins and 308 natural single-domain proteins (lengths 44 or 72 amino acids) together with experimentally measured unfolding free energies (Δ*G*). The fold-switch benchmark from Morpheus [18] classifies proteins as monostate or fold-switch based on published experimental evidence. The DIBS [20] catalogues intrinsically disordered regions that undergo coupled folding and binding upon interaction with a structured partner. The ATLAS database [21] provides three independent 100 ns all-atom molecular dynamics trajectories per protein for 1,290 proteins. The mean per-residue RMSF values across replicates served as the reference for per-residue validation. The OC23 and OC85 benchmarks [22] contain proteins crystallised in both open and closed conformations, stratified by TM-score between states below (OC23, *n* = 23) or above (OC85, *n* = 21) 0.85. The human phosphorylation dataset was assembled from PhosphoELM [23] as described below. The *Escherichia coli* K-12 reference proteome (UniProt UP000000625) was downloaded from UniProt; all proteins with sequence lengths between 50 and 2,000 amino acids were retained (*n* = 4,188).

### 7.2 Identification of buried phosphorylation proteins

Experimentally validated phosphorylation events on serine, threonine, and tyrosine residues were retrieved from PhosphoELM [23]. For each protein, per-residue solvent-accessible surface area (SASA) was computed on the corresponding crystal structure using the Rosetta software suite [32]. A phosphorylation site was classified as buried if its SASA was less than 20%. Proteins harbouring at least one buried phosphorylation site were retained as the positive set (*n* = 492).

### 7.3 Embedding extraction

All embeddings were extracted using the 360M-parameter model variant of SimpleFold [15]. The 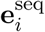 (sequence-conditioned) embedding was obtained by a single forward pass with all atomic coordinates set to **0**. The structural ensemble was generated by running the Euler–Maruyama sampler with 25 steps on a log-spaced timestep schedule and sampling temperature *τ* = 0.3, independently from *K* = 10 different Gaussian noise seeds. At the final denoising step, a per-residue latent tensor of shape [*L* × *D*] was recorded for each of the *K* conformations.

### 7.4 Score computation

For each protein, the PROTEUS score is the maximum per-residue Euclidean distance between the sequence-only embedding and the mean structural embedding:

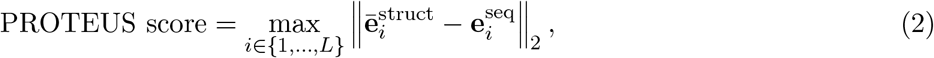

where 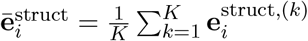 is the mean over the *K* = 10 structural embeddings at residue *i*. The fold-switch benchmark served as the development set for score selection. Approximately 70 candidate scoring functions derived from the same trajectory embeddings were evaluated by AUROC on this benchmark, including per-residue L2 norms, cosine distances, ensemble variance, and trajectory nonlinearity metrics. The score computed as above (l2 delta max) consistently ranked highest and was adopted as the primary score. All subsequent benchmarks, including the ATLAS MD fluctuation validation, open/closed state discrimination, and buried phosphorylation site detection, were evaluated after score selection was complete and are therefore fully independent of this development process.

For the continuum and ATLAS analyses, PROTEUS scores were length-residualised by regressing on sequence length via ordinary least squares and retaining the residuals, removing the dependence of the per-residue maximum on protein size. For the *E. coli* proteome analysis, proteins were filtered for structural order using AIUPred [19]: proteins with mean AIUPred score *>* 0.3 or with more than 30% of residues predicted as disordered (AIUPred score ≥ 0.3) were excluded before ranking.

### 7.5 Statistical analysis

Area under the receiver-operating characteristic curve (AUROC) was computed using scikit-learn. 95% confidence intervals were estimated by non-parametric bootstrap with 1,000 replicates using the percentile method. Group differences were assessed with the two-sided Mann–Whitney *U* test. Partial Spearman correlations were computed by regressing both the PROTEUS score and the response variable against the confounding variables (sequence length and, where indicated, pLDDT or crystallographic resolution) via ordinary least squares, then computing Spearman *ρ* on the residuals. Within-protein Spearman *ρ* between per-residue PROTEUS profiles and MD RMSF was computed independently for each protein in the ATLAS dataset. The median over all proteins is reported as the summary statistic.

### 7.6 Data and code availability

All code for embedding extraction, score computation, and statistical analysis is available at https://github.com/lupiochi/proteus.

