## Supplementary Information for "Unraveling protein conformational plasticity with PROTEUS"

### Supplementary Information for
### *Unraveling protein conformational plasticity with PROTEUS*

Luiz Felipe Piochi[1], Yasaman Karami[1,*], and Hamed Khakzad[1,*]

[1]Université de Lorraine, CNRS, Inria, LORIA, F-54000 Nancy, France

#### Contents

**S1 Supplementary Tables** **2**

**S2 Supplementary Figures** **3**

### S1 Supplementary Tables

Table S1: **Composition of datasets used in this study.** Counts are the number of proteins retained for analysis after filtering. Length range is given in amino acids. "Reference" indicates the original publication or database from which the dataset was obtained.

| Dataset | Class / role | $n$ | Length (aa) | Reference |
|---|---|---|---|---|
| Tsuboyama *de novo* | rigid scaffolds (positive control) | 146 | 44, 72 | [1] |
| Tsuboyama natural | natural single-domain proteins | 308 | 44, 72 | [1] |
| Morpheus monostate | single-fold proteins (negative class) | 197 | 17–510 | [2] |
| Morpheus fold-switch | fold-switching proteins (positive class) | 189 | 17–510 | [2] |
| DIBS | folding-upon-binding IDPs | 736 | 14–500 | [3] |
| ATLAS (full) | MD reference ($3\times100$ ns/protein) | 1,290 | 50–800 | [4] |
| ATLAS confound subset | stratified for pLDDT/AIUPred control | 300 | 50–800 | this study |
| ATLAS rigid | MD-validated rigid (mean RMSF $\leq 1$ Å) | 380 | 50–800 | this study |
| OC23 | open/closed pairs, TM-score $< 0.85$ | 23 | 90–540 | [5] |
| OC85 | open/closed pairs, TM-score $> 0.85$ | 21 | 90–540 | [5] |
| PhosphoELM buried sites | buried phospho-sites (SASA $< 20\%$) | 492 | 80–2000 | [6] |
| *E. coli* K-12 (UP000000625) | full reference proteome | 4,188 | 50–2000 | UniProt |
| *E. coli* ordered subset | post AIUPred filter ($\leq$0.3 mean and frac.) | 3,549 | 50–2000 | this study |

Table S2: **SimpleFold inference hyperparameters used for PROTEUS scoring.** All embeddings extracted with the 360M parameter SimpleFold variant. Sampler settings were used uniformly across all datasets in this study.

| Parameter | Value |
|---|---|
| Model variant | SimpleFold-360M |
| Sampler | Euler–Maruyama |
| Number of sampling steps | 25 |
| Timestep schedule | log-spaced, $t \in [0, 1]$ |
| Sampling temperature $\tau$ | 0.3 |
| Conformations per protein $K$ | 10 |
| Latent embedding dim. $D$ | 1024 |
| Sequence-only regime | coordinates = $\mathbf{0}$, single forward pass at $t = 0$ |
| Structural regime | per-residue mean over $K$ samples at $t = 1$ |

#### S2 Supplementary Figures

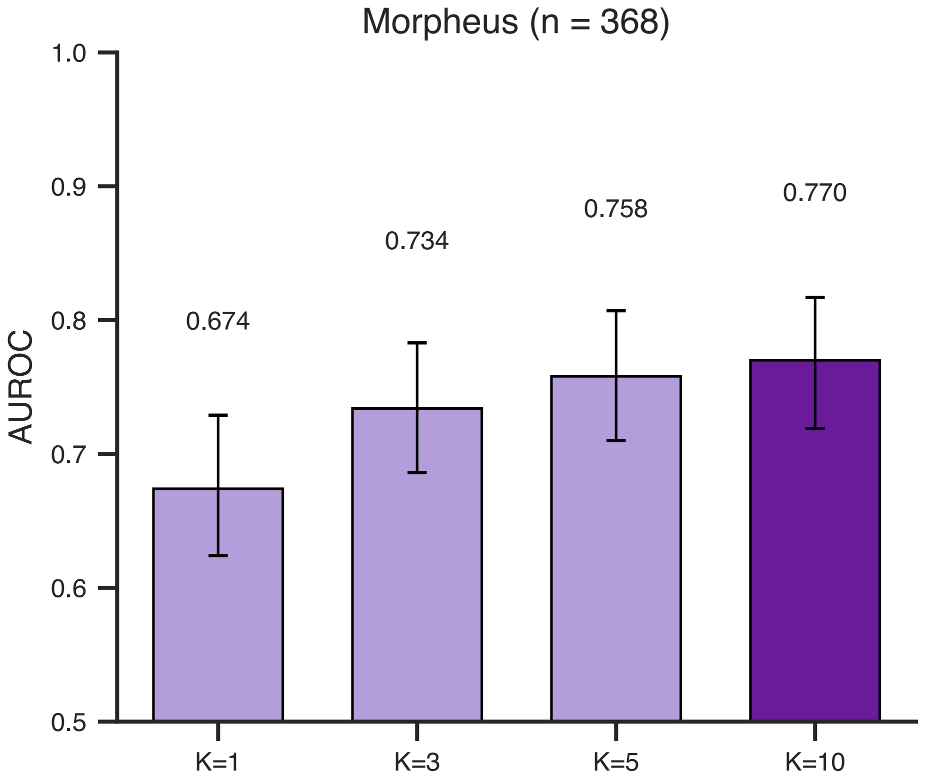


Figure S1: **Effect of ensemble size $K$ on PROTEUS discrimination.** AUROC for fold-switch versus monostate classification on the Morpheus benchmark ($n = 386$) as a function of the number of independent denoising trajectories $K$ used to construct the mean structural embedding $\bar{\mathbf{e}}_i^{\text{struct}}$. AUROC increases monotonically from 0.674 at $K = 1$ to 0.770 at $K = 10$, with diminishing returns beyond $K = 5$. Error bars are bootstrap 95% confidence intervals (1,000 replicates). $K = 10$ was adopted as the default for all subsequent analyses.

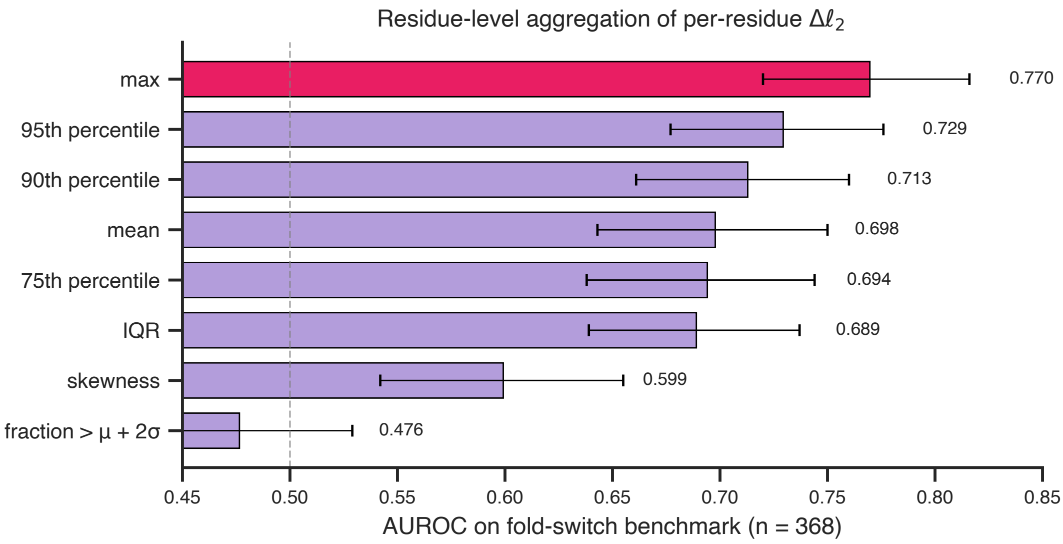


Figure S2: **Residue-level aggregation of the per-residue $\Delta\ell_2$ vector.** AUROC on the Morpheus fold-switch benchmark ($n = 386$) for eight alternative aggregators of the per-residue Euclidean displacement between sequence-only and mean structural embeddings: maximum (red), 95th percentile, 90th percentile, mean, 75th percentile, IQR, skewness, and the fraction of residues exceeding $\mu + 2\sigma$. The maximum residue achieves the highest AUROC (0.770), confirming that the most ambiguous residue carries the discriminative signal whereas global summaries dilute it. Error bars are bootstrap 95% confidence intervals.

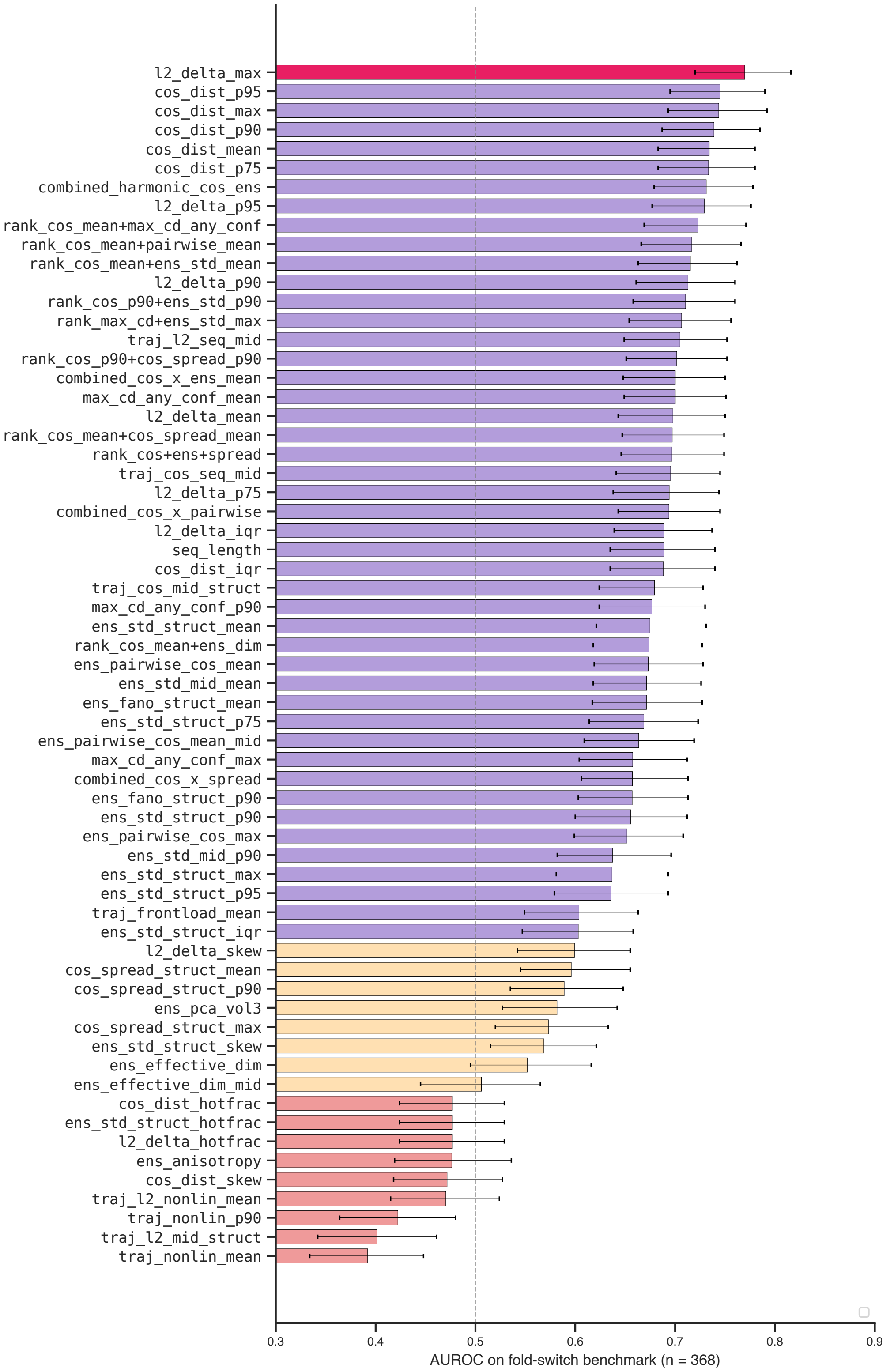


Figure S3: **Comprehensive score-selection screen on the fold-switch benchmark.** Approximately 70 candidate scoring functions derived from sequence-only and structural embeddings were evaluated by AUROC ($n = 386$). Candidates include per-residue $\ell_2$ displacement statistics ($\ell_2$_delta_*), cosine distances (cos_dist_*), ensemble spread and Fano factors (ens_std_*, ens_fano_*), pairwise cosine summaries (ens_pairwise_*), trajectory non-linearity (traj_nonlin_*, traj_l2_mid_struct), ensemble effective dimensionality, and rank-based combinations. l2_delta_max (red) ranks first. Trajectory non-linearity scores (bottom, pink) perform at or below chance, indicating that the discriminative signal is concentrated at the trajectory endpoints rather than in mid-trajectory curvature. Error bars are bootstrap 95% confidence intervals.

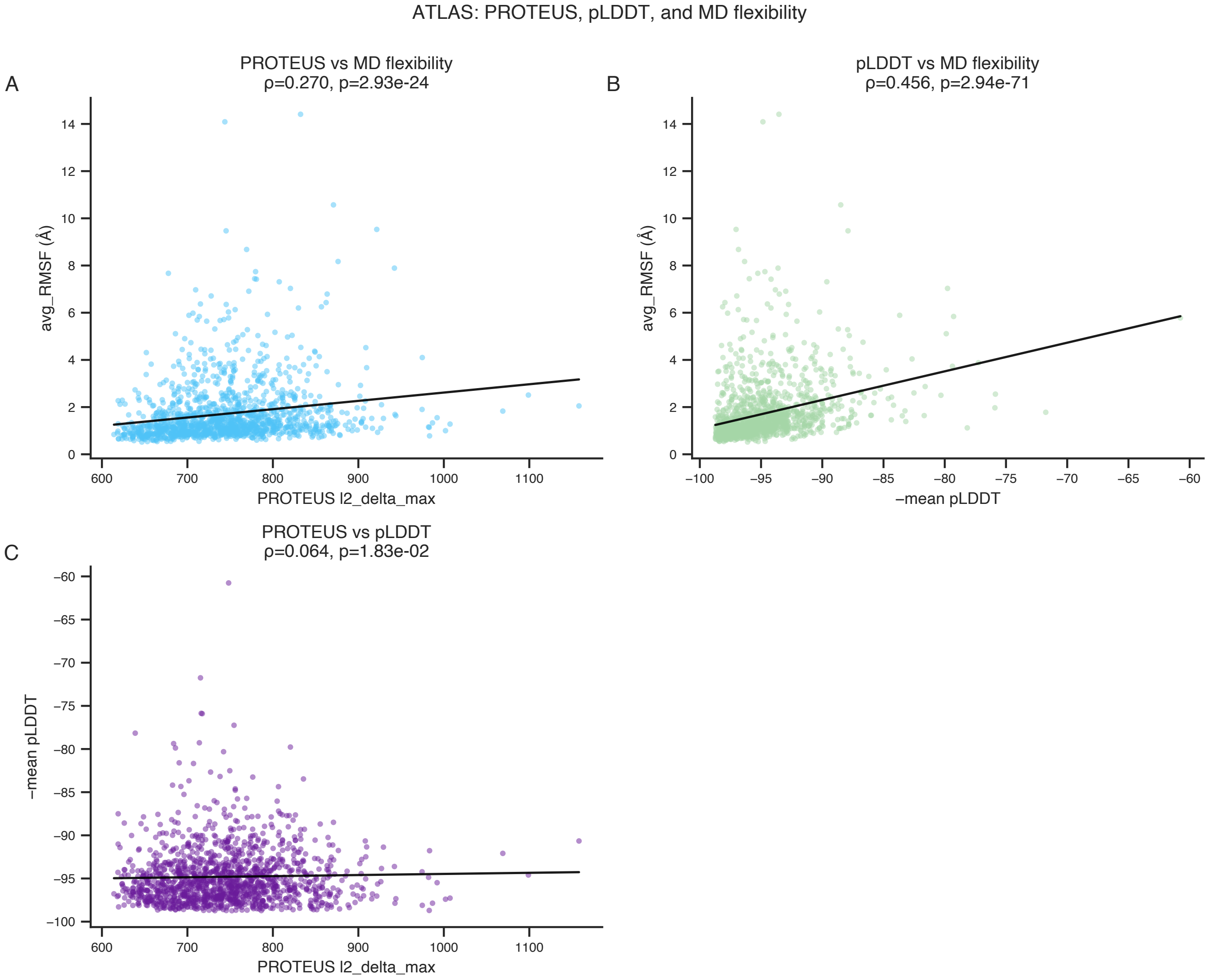


Figure S4: **Pairwise associations between PROTEUS, pLDDT, and MD flexibility on ATLAS ($n$ = 1,290).** **A**, Protein-level PROTEUS (l2_delta_max) versus mean MD RMSF (Spearman $\rho = 0.270$, $p = 2.93 \times 10^{-24}$). **B**, Mean pLDDT (negated, so increasing $x$ corresponds to lower confidence) versus mean MD RMSF ($\rho = 0.456$, $p = 2.94 \times 10^{-71}$): pLDDT is the stronger univariate predictor of average flexibility. **C**, PROTEUS versus mean pLDDT ($\rho = 0.064$, $p = 1.83 \times 10^{-2}$): the two scores are nearly orthogonal at the protein level, supporting their use as complementary predictors of conformational plasticity. Solid lines are OLS fits.

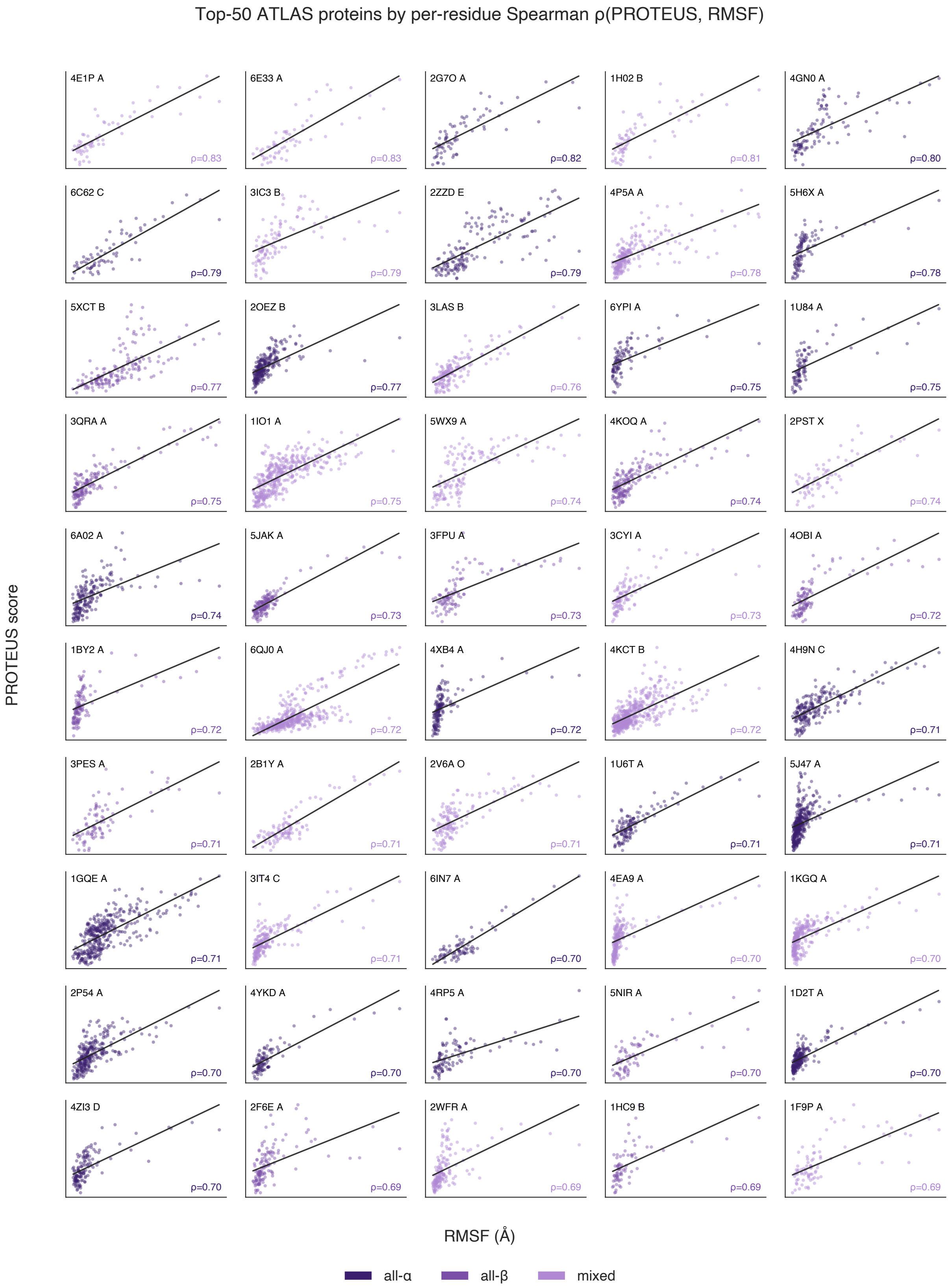


Figure S5: **Per-residue PROTEUS–RMSF concordance for the top-50 ATLAS proteins.** Scatter plots of per-residue PROTEUS score versus mean MD RMSF for the 50 ATLAS proteins with the highest within-protein Spearman $\rho$. Each dot represents one residue; the within-protein $\rho$ is annotated in the lower-right corner of each panel. Colour encodes SCOP secondary-structure class: dark purple (all-$\alpha$), medium purple (all-$\beta$), and light purple (mixed). Solid black lines are OLS fits.

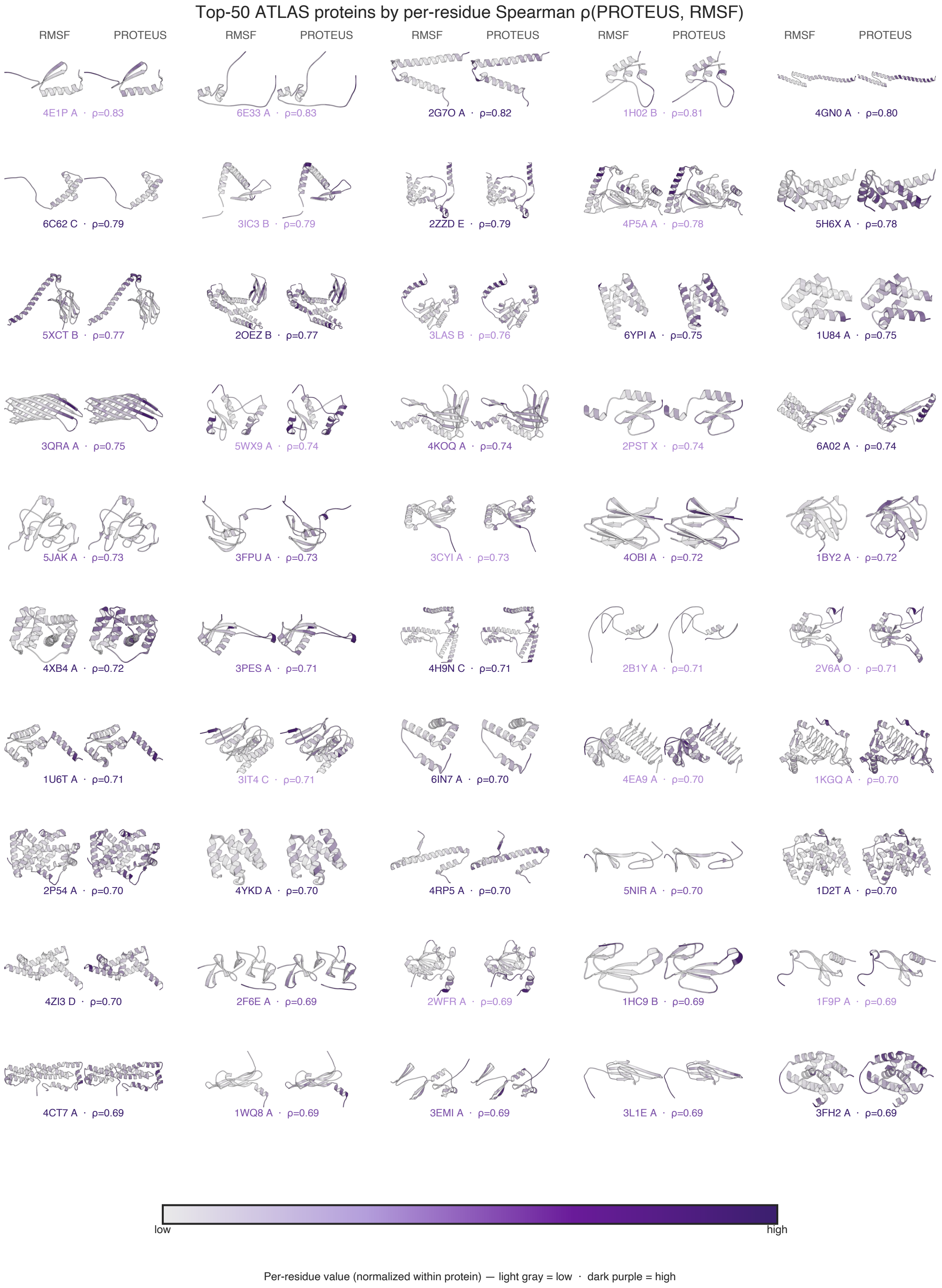


Figure S6: **Structural visualisation of the top-50 ATLAS proteins.** Paired cartoon representations of the same 50 proteins shown in Figure S5, coloured by mean per-residue MD RMSF (left of each pair) and per-residue PROTEUS score (right of each pair). Both scales are min–max normalised within each protein (light grey = low, dark purple = high). Within-protein Spearman $\rho$ is annotated next to each PDB code. Spatial concordance between the two colourings illustrates that PROTEUS identifies the same flexible regions as molecular dynamics across $\alpha$, $\beta$, and mixed folds.

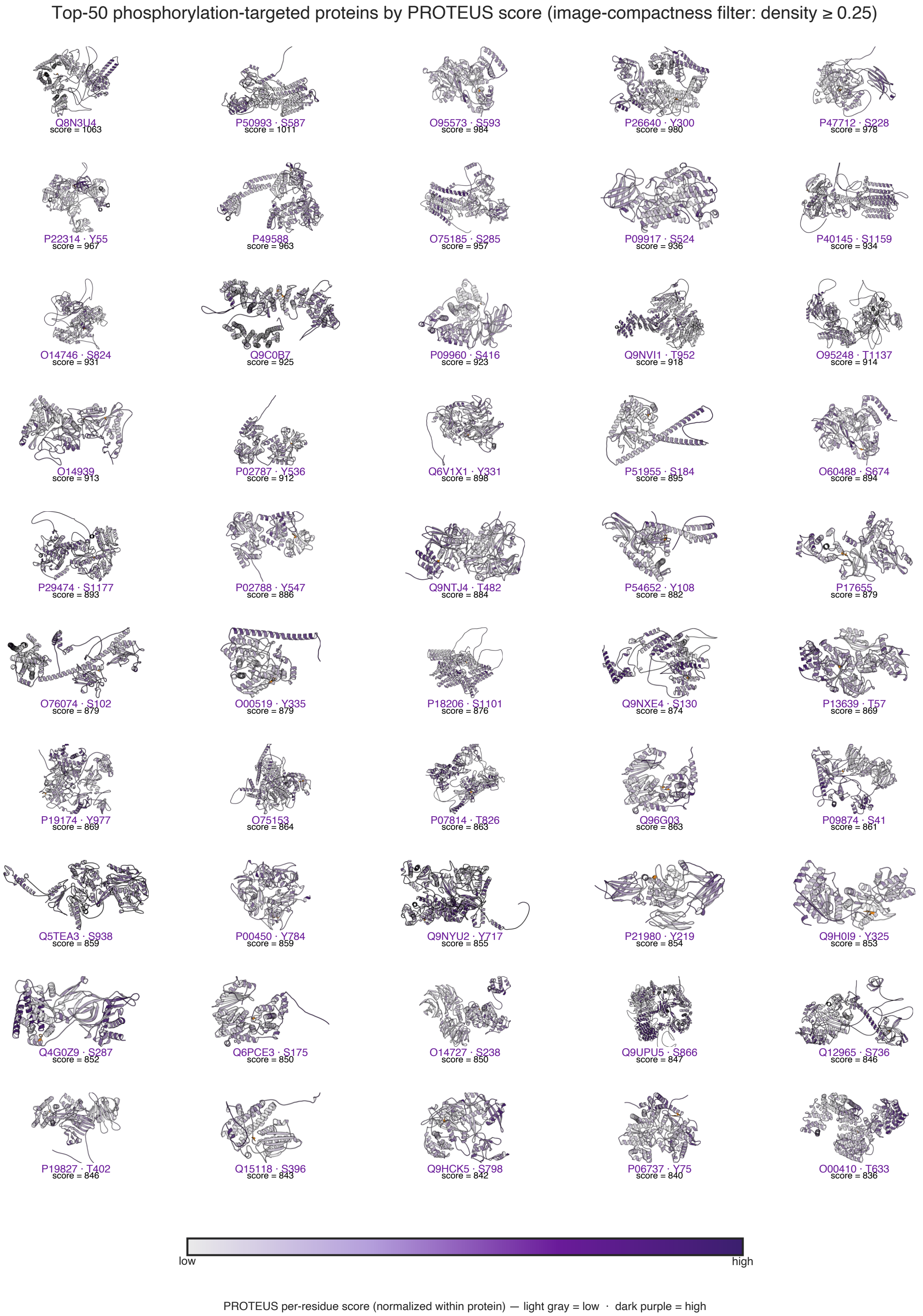


Figure S7: **Top-50 PROTEUS-scoring proteins from the buried-phosphorylation-site dataset.** Cartoon representations coloured by per-residue PROTEUS score (min–max normalised within each protein; light grey = low, dark purple = high). Each panel is annotated with the UniProt accession of the host protein, the phosphorylation site (residue type and position), and the protein-level PROTEUS score. The ranking is computed on the full dataset ($n = 492$).

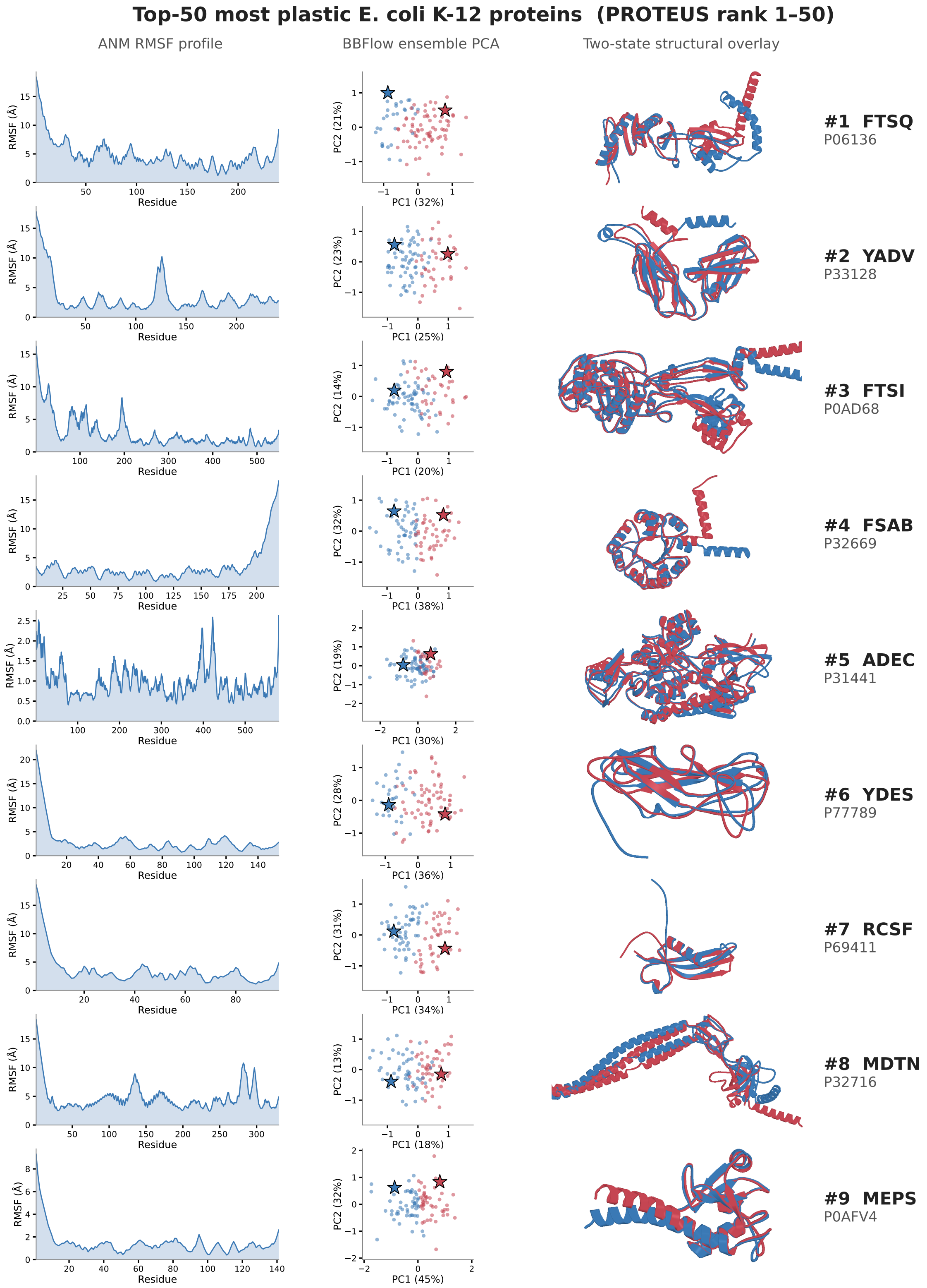


Figure S8: **BBFlow ensemble characterisation of the top-50 most plastic *E. coli* K-12 proteins (PROTEUS rank 1–50).** For each protein, three panels are shown side-by-side: *left*, anisotropic network model (ANM) RMSF profile per residue (Å); *centre*, PCA projection of pairwise RMSD between BBFlow-sampled conformations (each dot is one sample; the blue and red stars indicate two representative conformations selected from the PC1/PC2 extremes); *right*, two-state structural overlay of the blue and red representative conformations. UniProt accession and gene symbol are annotated to the right of each row. Page 1 shows ranks #1–9; pages 2–6 continue ranks #10–50.

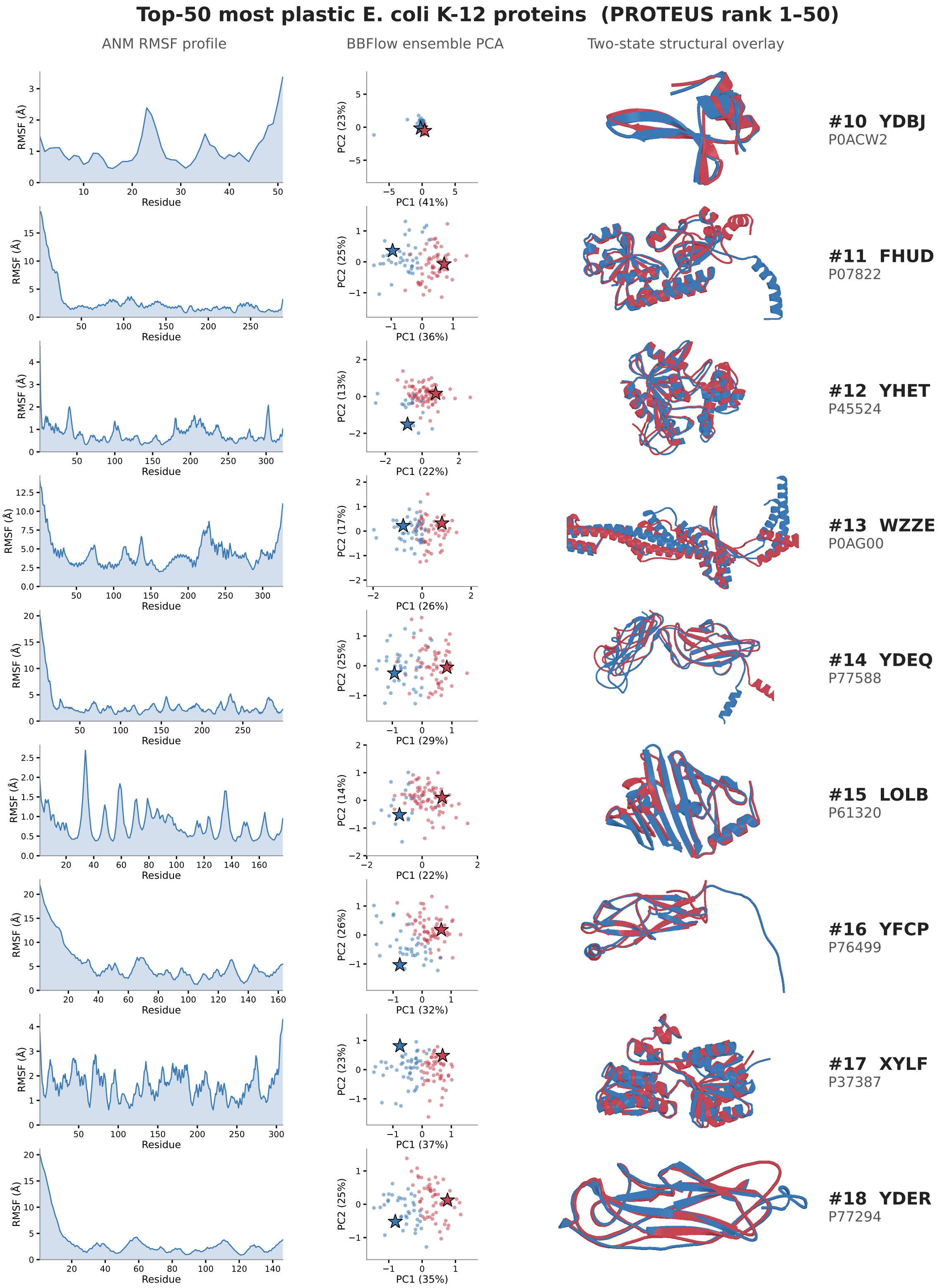


**Figure S8 (continued).** Top-50 *E. coli* K-12 plastic proteins, ranks #10–18.

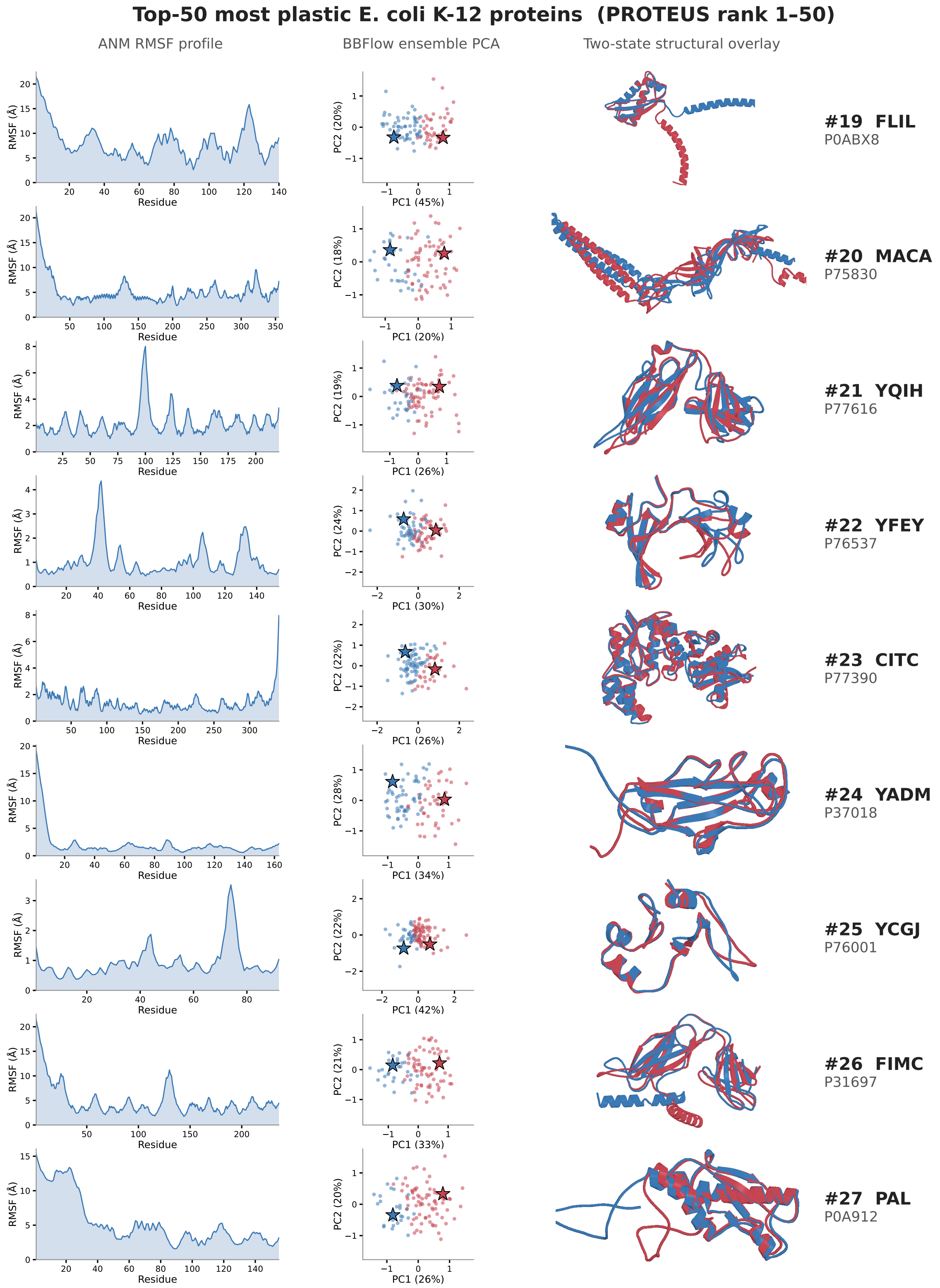


**Figure S8 (continued).** Top-50 *E. coli* K-12 plastic proteins, ranks #19–27.

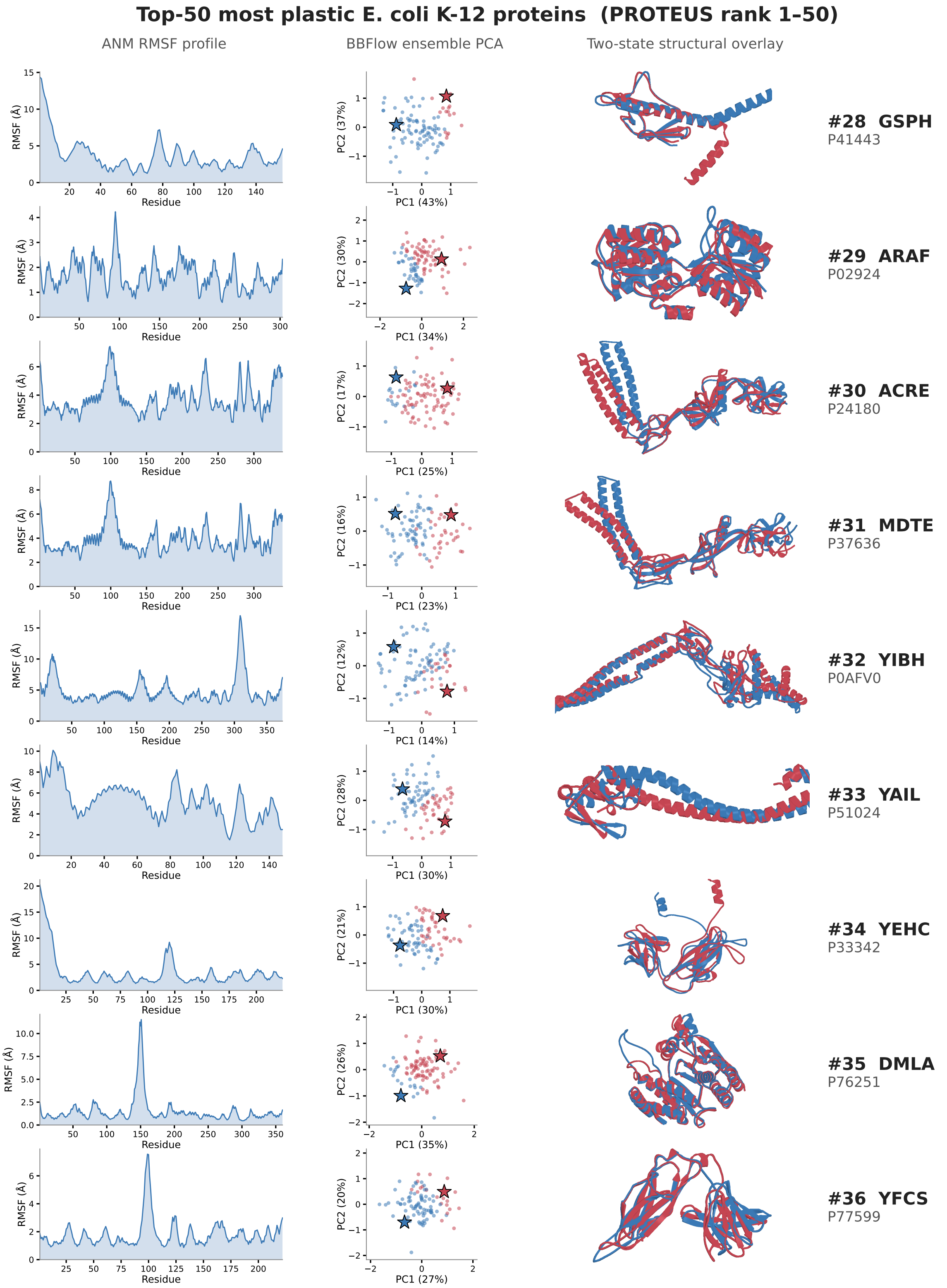


**Figure S8 (continued).** Top-50 *E. coli* K-12 plastic proteins, ranks #28–36.

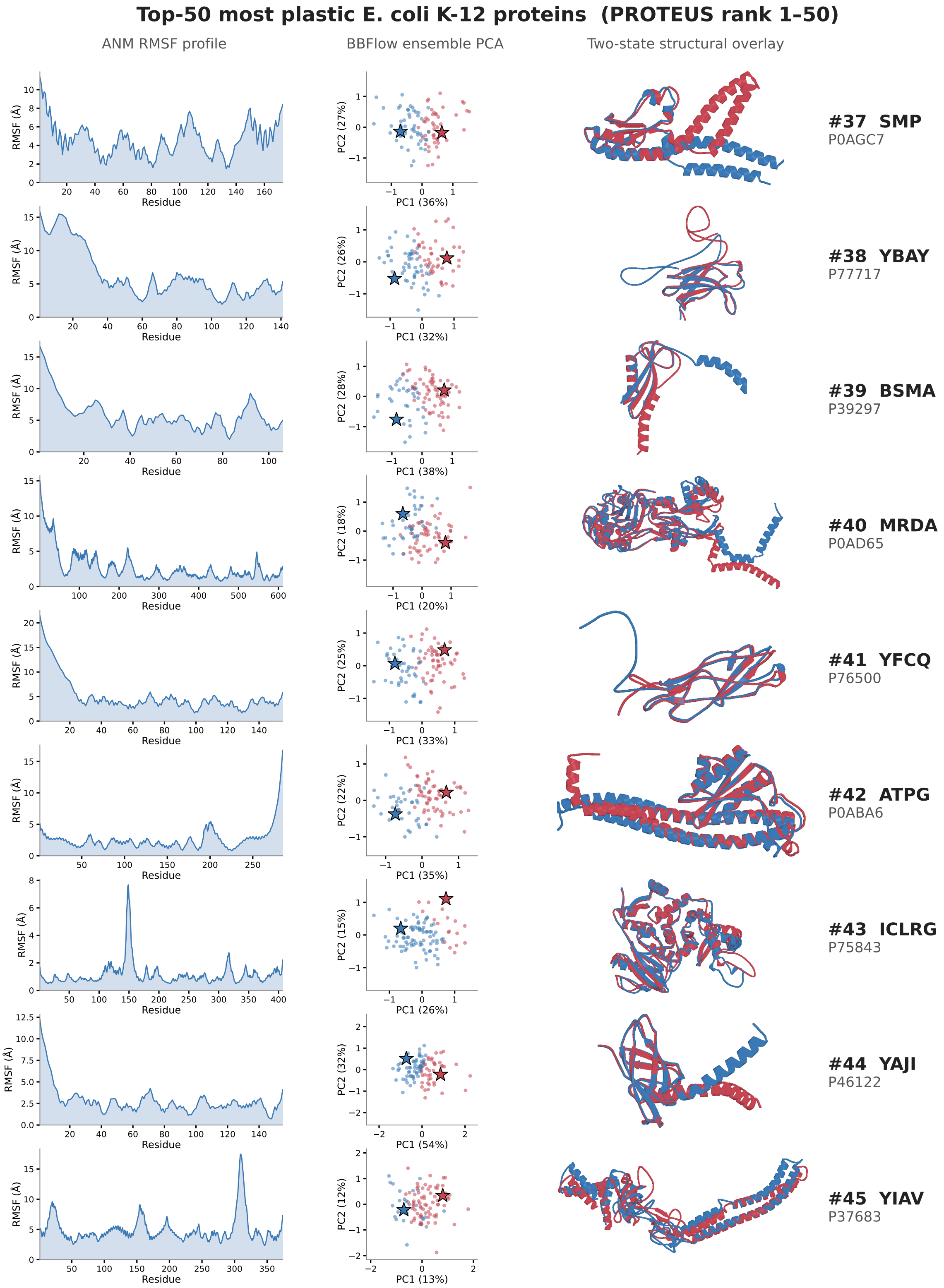


**Figure S8 (continued).** Top-50 *E. coli* K-12 plastic proteins, ranks #37–45.

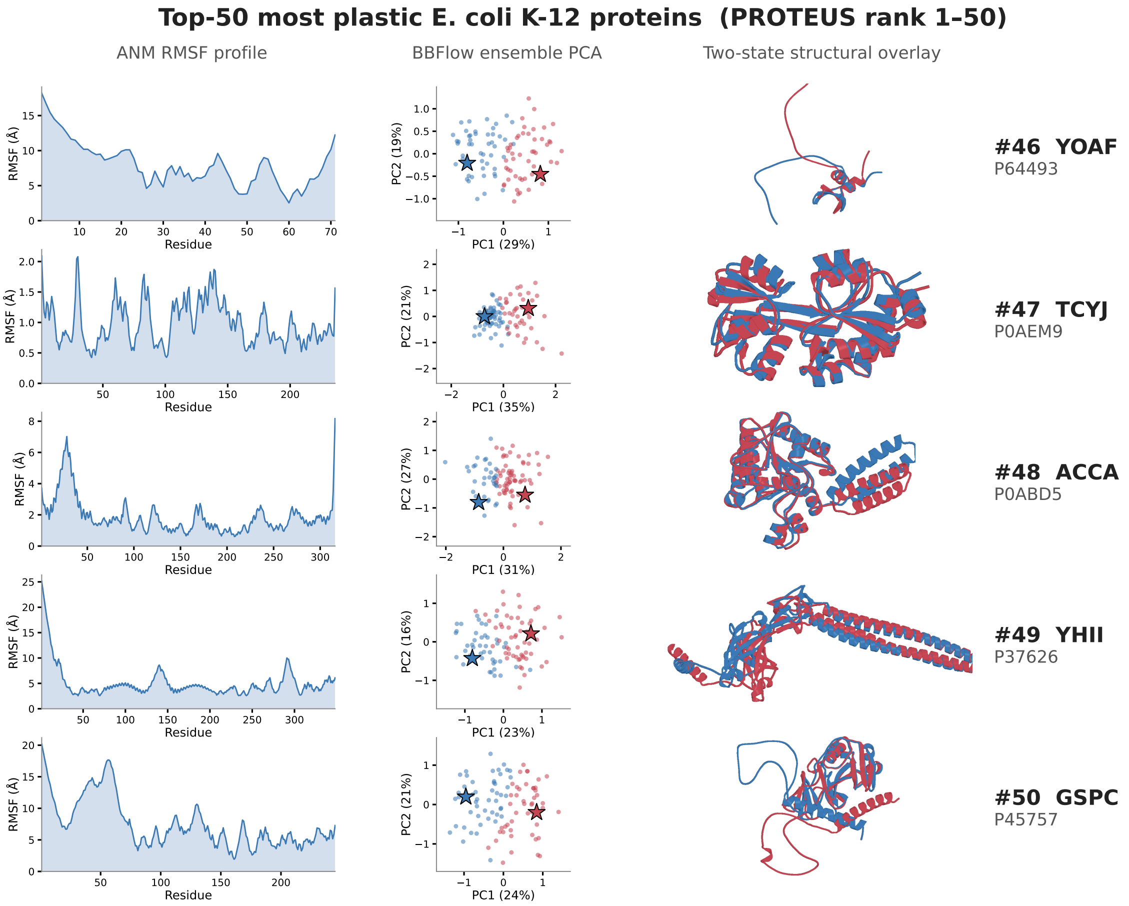


**Figure S8 (continued).** Top-50 *E. coli* K-12 plastic proteins, ranks #46–50.

4. Yann Vander Meersche, Gabriel Cretin, Aria Gheeraert, Jean-Christophe Gelly, and Tatiana Galochkina. ATLAS: protein flexibility description from atomistic molecular dynamics simulations. *Nucleic Acids Research*, 52(D1):D384–D392, January 2024.

5. Yogesh Kalakoti and Björn Wallner. AFsample2 predicts multiple conformations and ensembles with AlphaFold2. *Communications Biology*, 8(1):373, March 2025.

6. Holger Dinkel, Claudia Chica, Allegra Via, Cathryn M. Gould, Lars J. Jensen, Toby J. Gibson, and Francesca Diella. Phospho.ELM: a database of phosphorylation sites—update 2011. *Nucleic Acids Research*, 39(suppl_1):D261–D267, January 2011.
